# Revisiting regulatory decoherence and phenotypic integration: accounting for temporal bias in co-expression analyses

**DOI:** 10.1101/2021.04.08.438389

**Authors:** Haoran Cai, David L. Des Marais

## Abstract

Environment can alter the degree of phenotypic variation and covariation, potentially influencing evolutionary trajectories. However, environment-driven changes in phenotypic variation remain understudied. In an effort to exploit the abundance of RNASequencing data now available, an increasing number of ecological studies rely on population-level correlation to characterize the plastic response of the entire transcriptome and to identify environmentally responsive molecular pathways. These studies are fundamentally interested in identifying groups of genes that respond in concert to environmental shifts. We show that population-level differential co-expression exhibits biases when capturing changes of regulatory activity and strength in rice plants responding to elevated temperature. One possible cause of this bias is regulatory saturation, the observation that detectable co-variance between a regulator and its target may be low as their transcript abundances are induced. This phenomenon appears to be particularly acute for rapid-onset environmental stressors. However, our results suggest that temporal correlations may be a reliable means to detect transient regulatory activity following rapid onset environmental perturbations such as temperature stress. Such temporal bias is likely to confound the studies of phenotypic integration, where high-order organismal traits are hypothesized to be more integrated with strong correlation under stressful conditions, while recent transcriptome studies exhibited weaker coexpression between genes under stressful conditions. Collectively, our results point to the need to account for the nuances of molecular interactions and the possibly confounding effects that these can introduce into conventional approaches to study transcriptome datasets.

## 1 INTRODUCTION

Organisms evolve to maximize their individual performance under dynamically changing environments. The capacity of a single genotype to generate a range of environmentally induced phenotypes is known as phenotypic plasticity. Phenotypic plasticity has been widely documented (Stotz et al. 2021) and has recently received considerable attention as it may allow populations to persist in the face of rapid climate change (West-Eberhard 2003, Oostra et al. 2018, Nicotra et al. 2010, Gibert et al. 2019). Understanding the molecular and genetic basis of phenotypic plasticity has long been a goal for evolutionary genetics (Smith 1990, Bradshaw 1965) as a means to dissect trait co-variances and to predict the functional consequences of variable plastic mechanisms within and between populations (Des Marais et al. 2013, Palakurty et al. 2018, Tanner et al. 2022). Gene expression neatly bridges an organism’s genotype to its cellular biology and, by extension, higher-order developmental and physiological processes (Wray et al. 2003, Carroll 2008) and, at genome scale, is orchestrated by the underlying “Transcription Regulatory Network” (TRN) (Gibson 2008). Transcription factors (TF) are key nodes in TRNs that regulate the expression of other genes (Buchanan et al. 2010), coordinating the entire transcriptional program, and are often found to drive large-effect loci associated with the environmental response (Alonso-Blanco et al. 2005, Yano et al. 2000, Fukao et al. 2011). Transcriptional regulation can be highly responsive to both external and internal environmental cues, although formally linking these cellular phenotypes to whole organism environmental responses remains a challenge.

Considering each gene’s expression separately may fail to capture the full picture of the evolution and plastic response of the transcriptome collectively. Examining gene modules with coherent expression will make it more tractable to identify groups of genes associated with ecologically relevant outcomes (Palakurty et al. 2018). Therefore, gene co-expression and network analysis have been widely used to study multiple genes together. For example, co-expression and clustering analysis have been used to understand how the environment alters the expression and function of suits of genes simultaneously (Yan et al. 2019, Lea et al. 2019, Tanner et al. 2022, Zhao et al. 2016, Wang et al. 2013, Palakurty et al. 2018, Schneider et al. 2014, Fu et al. 2014), or to identify evolutionarily conserved functional modules between species (Ferrari et al. 2018, Gao et al. 2012, Monaco et al. 2015, Ruprecht et al. 2017*a,b*, Horn et al. 2016). However, the ease of performing differential co-expression analyses using existing approaches such as WGCNA (Langfelder and Horvath 2008) can obscure the assumptions they make about regulatory interactions. It is critical to understand possible biases and confounding factors using co-expression and module analysis.

Phenotypic integration refers to the magnitude of correlations among groups of related traits in a given organism (Pigliucci 2003). Phenotypic integration and modularity represent important factors and constraints that influence the phenotypic plasticity and evolutionary trajectory (Villamil 2018, Gianoli and Palacio-López 2009, Pigliucci and Preston 2004, Schlichting 1989). The ecological significance of patterns of changing phenotypic integration is not fully understood. Previous studies in phenotypic integration hypothesized that the number and strength of significant correlations among traits increase with environmental stress (Waitt and Levin 1993, Schlichting 1989, Gianoli 2004, García-Verdugo et al. 2009, Chapin 1991). However, recent studies on molecular traits (e.g., genome-wide gene expression) provided seemlingly conflicting evidence: the degree of covariation tends to be higher under benign conditions than under stressful conditions (Lea et al. 2019, Tanner et al. 2022, Southworth et al. 2009, Anglani et al. 2014). We here hypotheze that low population-level trait-trait correlations in gene expression under a stressful condition may be caused by a widely observed phenomenon called regulatory saturation. Regulatory saturation can occur when the transcript abundance of regulators is too high such that extra transcript will not lead to increased transcription of target genes. In a regulatory saturation regime, a low population correlation does not necessarily mean that the regulatory activity between two genes is low. Conversely, it means the regulation and integration strength of two genes are tight and strong. In the present work, we account for such potential confounding factors by considering the temporal context of gene expression.

Extensive efforts have been made to exploit the so-called “fourth dimension” of environmental response — time — to better understand the dynamics of TRNs and to identify putative signaling pathways or key regulatory genes (Bechtold et al. 2016, Yeung et al. 2018, Varala et al. 2018, Zander et al. 2020, Song et al. 2016, Greenham et al. 2017, Windram et al. 2012, Gargouri et al. 2015, Alvarez-Fernandez et al. 2020). Here, we exploit this temporal component of gene co-expression to characterize the dynamic organismal response to environmental conditions using an existing data set in rice (Wilkins et al. 2016). We contrast two broad approaches for using correlations between the transcript abundances of two genes: temporal correlations and population correlations. Population correlations are correlations of multiple individual samples at a given time point. Temporal correlations are correlations of two transcripts’ abundance over a time course for an individual sample.

Our analyses and simulation demonstrate that multiple types of temporal bias can occur when analyzing gene expression data, any of which could confound ecological or evolutionary inference. First, the number, identity, and period between sampling time points may not capture the transient transcriptional response of genes and, by extension, their co-variance with other transcrips (Bar-Joseph et al. 2012). Second, population coexpression may fail to capture transient induction of regulatory activities when the gene-gene interaction is under a regulatory saturated regime. Regulatory saturation can be particularly misleading because a population correlation may capture the transient reponse of a single gene’s expression, yet miss responsive regulatory interactions if the duration of high regulatory activity is short. A potential implication of regulatory saturation points to the need to account for temporal bias when studying phenotypic integration, or the regulatory coherence for molecular traits. Our study provides evidence of increasing within-individual phenotypic integration following stress treatments. Third, temporal correlation can alleviate the second temporal bias and capture transient interactions. Collectively, our work emphasizes both that transient regulatory interactions may lead to bias in population transcriptomic analyses as well as offer an opportunity to understand the evolution of gene regulation in better detail if properly accounted for in analyses.

## 2 MATERIALS AND METHODS

### 2.1 Data retrieval

We utilized a TRN prior previously constructed by Wilkins et al. (2016), which was obtained from the integration of ATAC-seq (Assay for Transposase-Accessible Chromatin using sequencing) data and the CIS-BP database of TF binding motif (Weirauch et al. 2014). We elected not to use their complete “Environmental Gene Regulatory Influence Network” because the estimation of the final network relied on information from mRNA-seq time series data; the analyses presented here represent a unique approach to analyzing these data. Genes that had corresponding *cis*-regulatory elements of TF in a region of open chromatin in their promoter regions are identified as the target gene for a given TF (Wilkins et al. 2016). Note that the open accessible regulatory region derived from the ATAC-seq of rice leaves remained stable across multiple environmental conditions in the Wilkins et al. study. In total, this “network prior” has 38,137 interactions: 357 TFs were inferred to interact with 3240 target genes. Interactions can be between TFs and non-TF targets, or between two TFs.

The RNA-seq data derive from chamber-grown plants and were retrieved from the Gene Expression Omnibus (www.ncbi.nlm.nih.gov/geo/) under accession number GSE74793. This dataset comprises time-coursed libraries for four rice cultivars exposed to control (benign), heat shock, and water deficit conditions. Samples were collected with 15 min intervals for up to 4h for each of the treatments; specifically, 18 time points for controlled conditions; 9 time points for drought treatment; and 16 time points for heat treatment. Here we used a time window of nine time points in each condition for analysis. TF family annotations were downloaded from the Plant TF database (Riaño-Pachón et al. 2007), from which Heat Shock Factors are those with the TF family label “HSF”. “Known” drought-related TFs were obtained from https://funricegenes.github.io/ in June 2020 using the search terms “drought”, “ABA”, and “drought tolerance”.

### 2.2 Dynamic correlations for regulator-target pairs

The expression relationship observed between genes in a time series sample may be caused by the time lag inherent in molecular interactions, in this case, transcriptional regulation. Such time lag reflects the time required for a TF’s activity to influence the expression of its target genes because transcription and translation take place over non-negligible time periods. Traditional correlation coefficients (e.g., Pearson correlation) cannot account for the staggered relationship between a regulator and a target. Here, we used a metric we call Max Cross Correlation (MCC), building on the cross-correlation between transcript abundances estimated by RNA-Seq, to examine the activities of regulatory interactions. The MCC over the time course has a direction constraint (from regulator to target) to evaluate the regulatory status. Consider two discrete time series denoting *f*(*t*) (regulator) and *g*(*t*) (target), both of length of *N* number of time points, the cross-correlation function is defined as:

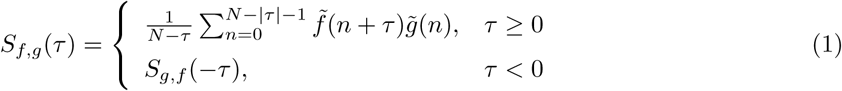

where the 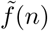 is a normalized time series (zero mean, unit variance). The maximum cross correlation *S*_*f,g*_(*τ*) is calculated under condition of *m* ≤ τ ≥ 0, where *m* is the max delay. The time delay that possesses the max correlation is defined as *τ*_*reg*_, representing the approximate time delay that occurs between a given regulatory-target pair. The max delay is set as 1, which in the current dataset represents a 15 min time interval. For comparison between maximum cross correlation distributions under multiple conditions, we used a Kolmogorov-Smirnov test using the *ks*.*test* function in *R*. Note that it is unknown to what extent of the temporal resolution the present method is effective in revealing the transient dynamic of the regulatory activity (the sampling interval in our dataset is 15 min).

### 2.3 Simulations for the minimal activation model

To illustrate the potential bias in capturing changing regulatory activities by using population level correlation, we implement simulation through a minimal activation model. The rate of production of TF *X* and gene *Y* (Fig. 4A) is described by the following equation:

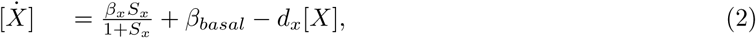

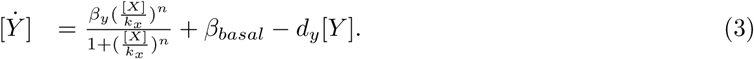

[*X*] and [*Y*] denote the mRNA concentrations of TF *X* and gene *Y* respectively. TF *X* affects the transcription of gene *Y*. The regulated expression of genes is represented by Hill function with cooperativity exponent *n*. It is assumed that each transcript degrades at a rate proportional to its own concentration (*d*_*x*_ and *d*_*y*_). Assume that the basal synthesis rate for *X* and *Y* is constant and equal with *β*_*basal*_. *β*_*y*_ can be taken as the maximum strength of regulations. The stochastic dynamics of the system are implemented through Gillespie stochastic simulation algorithm (Gillespie 1977). The Hill function can be tuned by the binding affinity *K*_*x*_ and the Hill coefficient *n*. The Hill coefficient quantifies the cooperativity of transcription factor bindings and thus the steepness of the sigmoidal stimulus-response curve of regulation. TFs often work cooperatively, where the binding of one TF to DNA enhances the binding of extra TFs. For example, a Hill coefficient *n >* 1 is indicative of positive cooperativity and thus, the system exhibits ultrasensitivity (Blüthgen et al. 2007). The other parameter, *K*_*x*_, indicates the binding affinity of a TF-DNA binding. We can manipulate the active range of regulatory interactions by these two parameters. A set of parameters including the induction signal strength *S*_*x*_ are determined to enable two regulatory regimes (Fig. 4C and 4F). Two types of perturbation imposed on cells at steady state are simulated, including press and pulse perturbations (Fig 4B). The press perturbation maintains the external signal at a certain high level throughout the time course, whereas the pulse perturbation indicates a discrete, transient induction of the external signal. We assume the external perturbation modulates the gene expression dynamics by the signal *S*_*x*_.

Temporal dynamics of TF *X* and gene *Y* were simulated for 100 times. The cross-correlation function is calculated for the bulk time series of *X* and *Y* (average of 100 simulations), whereas the population-level Pearson’s correlation coefficient (PCC) is calculated at each time point by using 100 simulations during the simulation.

## 3 RESULTS

### 3.1 Temporal bias in revealing dynamic regulatory interactions

We first evaluate the overall dynamic patterns of pairwise regulatory interactions by calculating the Maximum Cross Correlation (MCC, see Methods) for each pair of transcripts in a static network reported previously (Wilkins et al. 2016). The data comprise four rice cultivars grown under control, dehydration stress, or elevated temperature conditions; here we analyze the data by condition over a time duration of 135 minutes following the incidence of stress. Calculated MCC (Figure 1) is from all four cultivars were merged stress-wise. We set a threshold of 0.69 (p-value = 0.01) together with a fold-change cutoff (Fig. 2) according to the MCC under controlled conditions as the cutoff for the activation of regulatory interactions. We use the terms regulatory coherence and decoherence to mean increasing or decreasing co-expression, respectively. The coherence in our analyzes is reflected in higher MCCs under heat or dehydration stress conditions compared to control samples, as imposed by Wilkins et al.(Wilkins et al. 2016).

**Figure 1:**
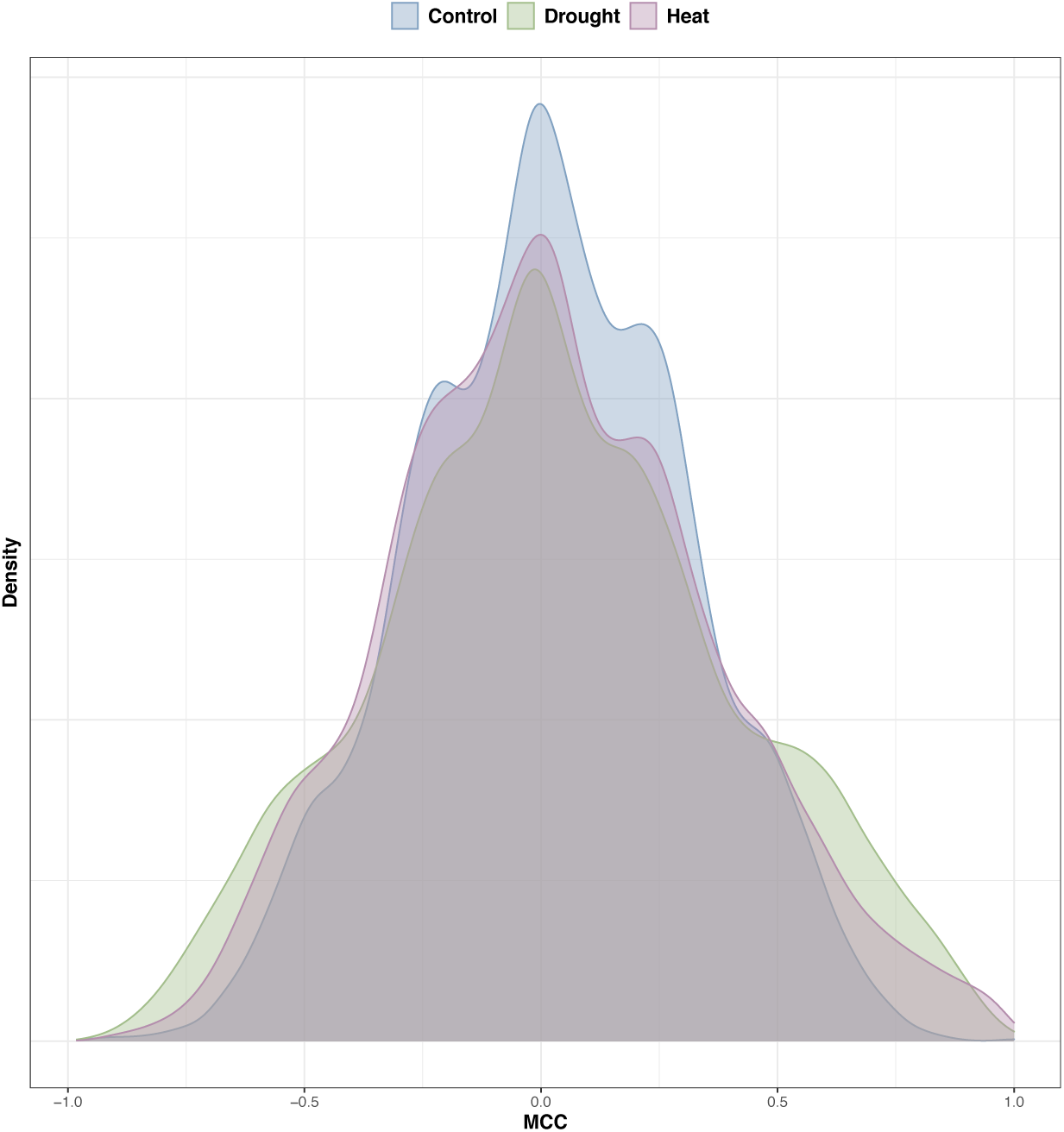
Temporal correlations under multiple environmental conditions show regulatory coherence. Temporal correlation is caculated as the Max Cross Correlation (with lag 1, see Methods) for each pair of transcripts, using a previously constructed static network prior. The data comprise four rice cultivars grown under control, dehydration stress, or elevated temperature conditions, and here we analyze the data by condition over a time duration of 135 minutes following the incidence of stress. Calculated Maximum Cross Correlations (MCC) from all four cultivars were merged stress-wise.

**Figure 2:**
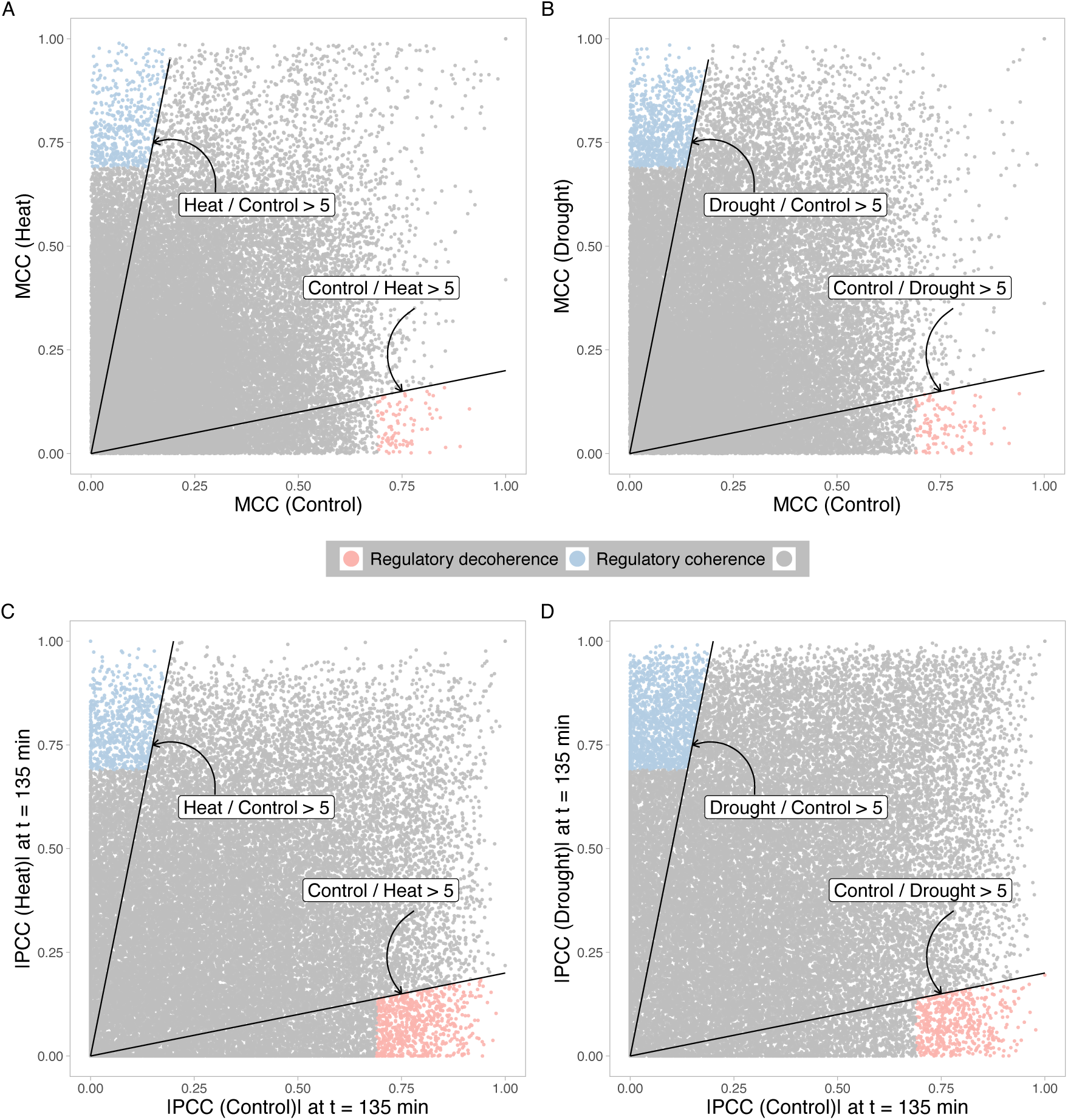
Environmental perturbations lead to contrasting patterns using temporal and population correlation. **A**. Comparison of the temporal correlation (Max Cross Correlation, MCC) for each regulator-target pair under control condition against heat condition. **B**. Comparison of the temporal correlation (MCC) for each regulator-target pair under control condition against soil drying condition. **C**. and **D**. show the Pearson Correlation Coefficient (PCC) of each regulator-target pair at 135 min after heat (**C**) and drought (**D**) treatment. The regulator-target pairs that are not significant in both conditions are in grey, for which the cutoff is 0.69 (p-value = 0.01). Red and blue labels highlight the pairs that show regulatory decoherence and regulatory coherence, respectively. Solid lines indicate that the ratio between regulatory scores under control and perturbed conditions is larger than 5.

The MCC distributions (Figure 1) reveal that stressful environments increase the coexpression strength among measured transcripts in the network prior. The distribution of the MCC under heat (KolmogorovSmirnov test statistic D = 0.0445, p-value < 2.2×10^−16^) and drought (Kolmogorov-Smirnov test statistic D = 0.0929, p-value < 2.2 × 10^−16^) conditions are significantly different from the control condition. We identified significant TF-gene interactions in stressful conditions which were not observed in the controlled condition, and vice versa. We found greater support for the former number: out of 38127 total interactions in the network prior, 496 and 839 pairs transition to active pairs in heat and drying stress, respectively (light blue points in Fig. 2A and 2B), whereas only 91 and 115 of them transition to inactive pairs under heat or drying, respectively. The observation of regulatory coherence is robust to various thresholds for activation (Fig. S1 and Fig. S2) and the max time lag (See Supporting Information A.2 and Fig. S3). Collectively, these results suggest a strong bias towards regulatory coherence in this rice expression dataset.

Our observation that co-expression increases with the onset of stress (regulatory coherence) is seemingly inconsistent with a recent study. Working with gene expression profiles of human monocytes exposed to a stress *in vitro*, Lea et al. calculated the differential population correlation among pairwise transcripts and found evidence supporting regulatory decoherence following perturbation (Lea et al. 2019). To explore the possible role of statistical methodology to explain the differing results of our two studies, we conducted a cross-sectional analysis by calculating population-level coexpression in the above rice gene expression data and our static network prior. Strikingly, for heat shock stress response, population correlations show little or no evidence of regulatory coherence under stress (Fig. 3A, S5 and S7). At many time points, the distributions of correlation coefficients under the stress condition are skewed towards regulatory decoherence (Fig. S5). These results are at odds with our observation of strong regulatory coherence in heat-stressed individuals when temporal correlations are assessed. An even more striking contrast is observed in the so-called heat shock regulon (Fig. 3A and Fig. 3B), formed by the Heat Shock Family TFs and their interactions with target genes in the static network prior, where the assessment using temporal correlations generates a starker contrast between control and stress condition than using population correlation at any time point. Such observation suggested that two approaches may capture different aspects of stress responses.

**Figure 3:**
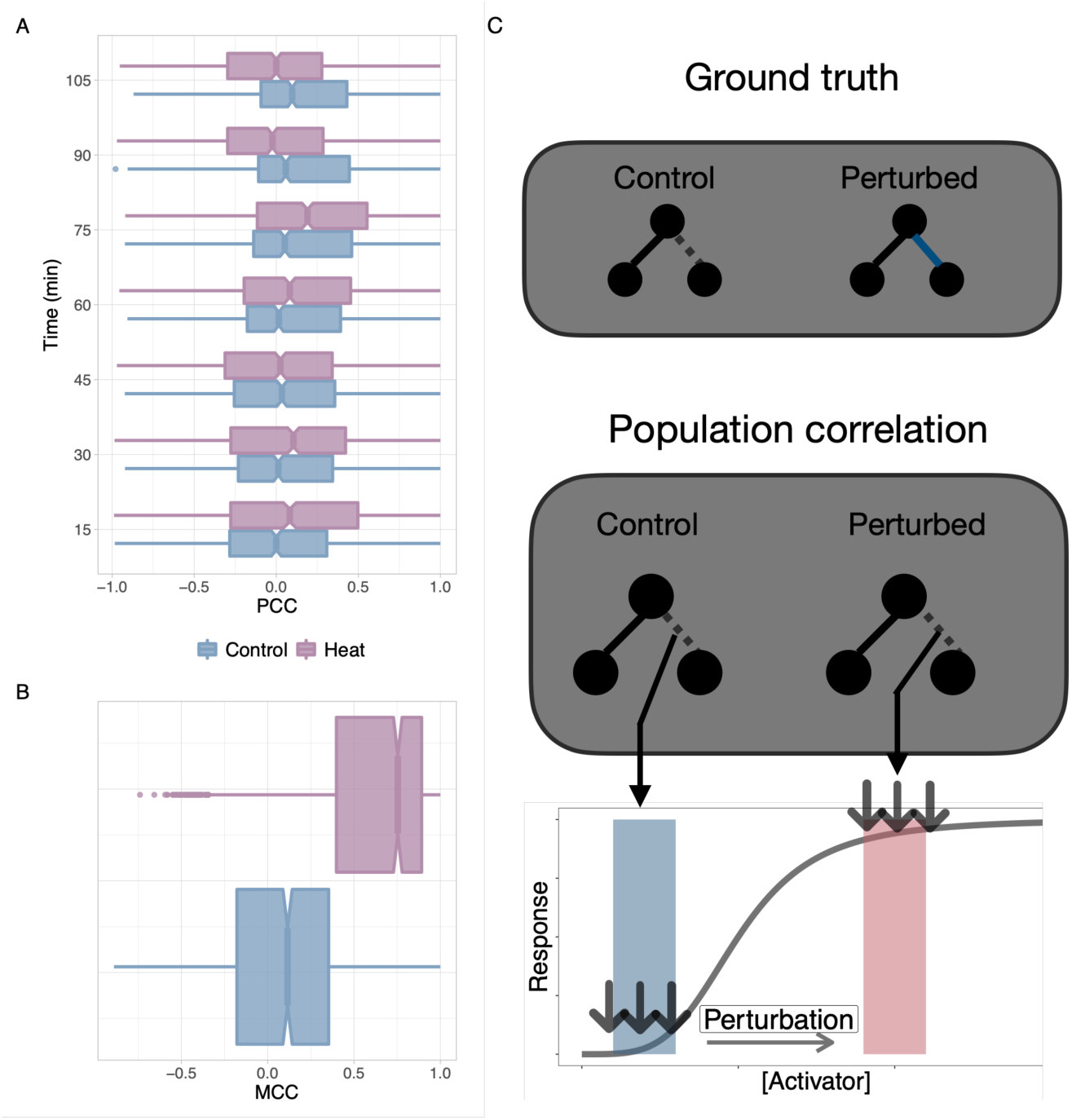
Heat shock regulon shows strong contrasting patterns upon heat shock treatment. **A.** Pearson Correlation Coefficient (PCC) under control and heat condition within heat shock regulon over the time course. Each boxplot represents the distribution of PCC under a given time and treatment. **B.** Max Cross Correlation (MCC) within the heat shock regulon. Genes in the heat shock regulon are identified by extracting links that include a regulator from the Heat Shock Family (HSF). As a family, HSFs have been demonstrated previously to show an important role in regulating genome-wide responses to elevated temperature in diverse species (Wang et al. 2004, Ohama et al. 2017). **C**. A schematic diagram depicts possible explanation of the temporal bias through regulatory saturation. The blue link is activated upon the perturbation (ground truth) by increasing the concentration of the regulator (an activator). However, if the dose-response curve is a sigmoid shape function, chances are the population correlation may not be able to detect such activation.

On the other hand, for drought stress response, population-based coexpression analysis shows regulatory coherence at several time points (Fig. S6). One confounding factor that may inflate estimated correlations, as pointed out by Lea et al. (2019), Parsana et al. (2019), is technical and latent biological covariates (e.g. genotype, cultivar effect) which may lead to spurious correlations. To explore how the inclusion of multiple genotypes in a population sample might affect correlation estimates, we use another dataset from *Brachypodium* with larger number of replicates for drought treatments across two inbred lines (Yun et al. 2021, in prep). In this larger, genotypically segregated sample, we still observe evidence of regulatory coherence in transcriptional response to drought treatment (Fig. S4).

### 3.2 Regulatory saturation as a cause for temporal bias

Through a simple mathematical model, we illustrate how regulatory saturation may be a confounding factor for identifying environmentally responsive transcriptional interactions. We contrast the outcomes of population-level metrics with our measure based on cross-correlation. A typical regulation function between a TF and a target gene (modeled as a dose-response curve) can be characterized as a Hill function (Alon 2019, Chu et al. 2009), which a is sigmoidal curve (See Methods) as shown in Fig. 4C and 4F (grey line). Two perturbation regimes are considered: A saturated regime in which additional TF transcripts beyond some concentration threshold fail to induce additive responses in their target genes, and a non-saturated regime characterized as the portion of a dose-response curve in which additional TF transcripts are associated with increased expression of their targets. We assume that the external perturbation modulates the gene expression dynamics by the signal *S*_*x*_. Smaller *K*_*x*_ and larger Hill coefficients *n*, indicative of higher binding affinity and cooperativity of transcription factor binding (See Methods), increase the probability of regulatory saturation after environmental perturbation. Saturation of the regulatory interaction effectively masks the differential regulatory interaction upon perturbation, even if the TF *X* is nominally an environmentally induced activator of the gene *Y*. In addition, two possible external perturbation regimes are simulated (Fig. 4B): press perturbation and pulse perturbation, which differ in the duration of the perturbation imposed on the given regulatory pair. If the upstream signal for a TF-gene pair has the property of adaptation, the signal induction may only last for a short period of time. Adaptation here is defined by the ability of circuits to respond to input change but to return to the pre-stimulus output level, even when the input change persists (Ma et al. 2009, Briat et al. 2016).

**Figure 4:**
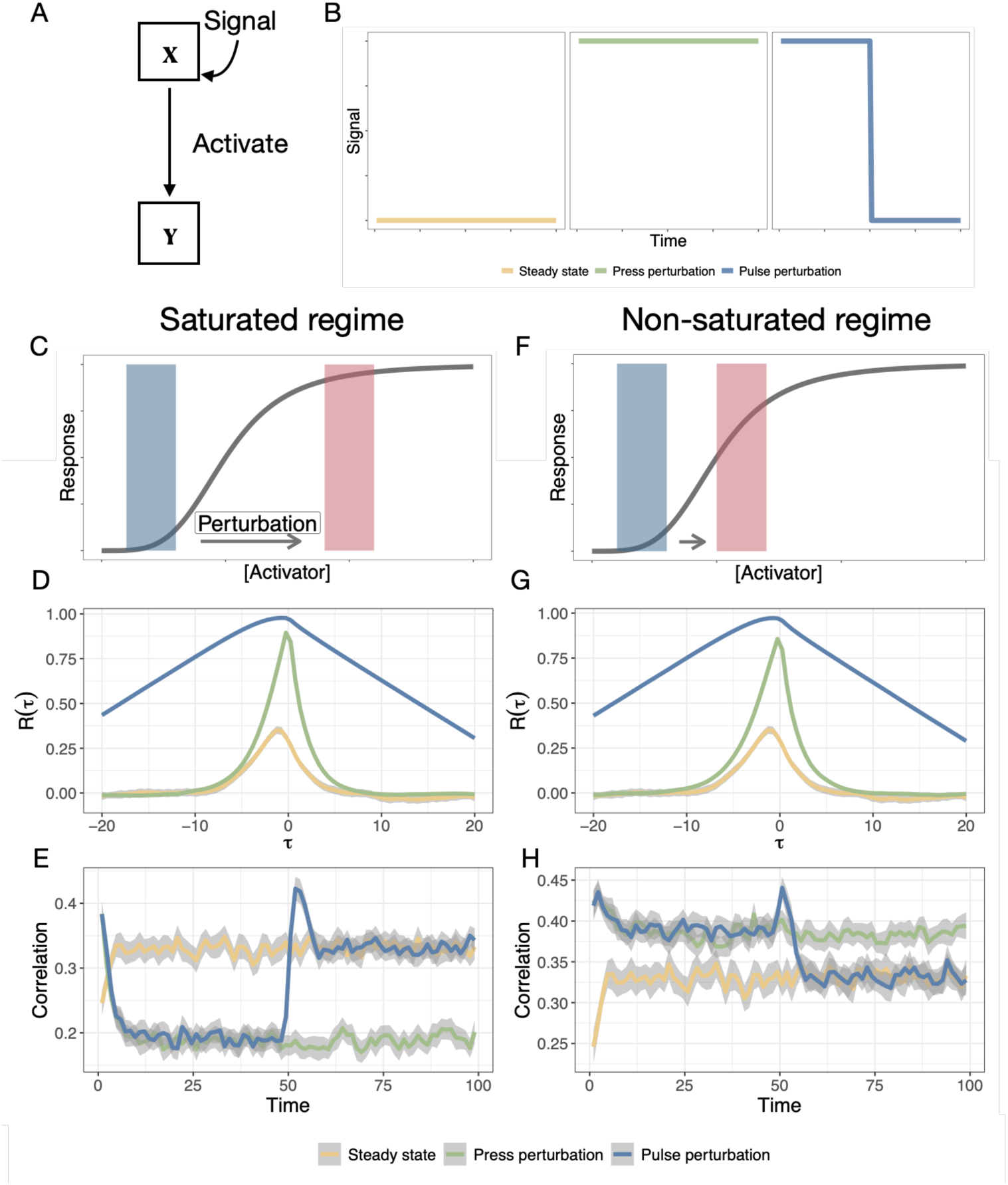
Illustrated examples through stochastic simulation indicate the robustness of using temporal correlation to detect regulatory coherence. The population level correlation may lead to temporal bias in detecting regulatory coherence depending on the regulatory regime. **A**. A schematic illustration of the minimal activation model explored here and **B**. input signals corresponding to three perturbation scenarios. **C - E**. The cross correlation function and population-level correlation between activator *X* and target *Y* under a saturated regime. The cross correlation function robustly reveals a peak in response to perturbations while the perturbation may lead to reduction of correlation when using population correlation over the time course. **F - H**. The cross correlation function and population-level correlation between activator *X* and target *Y* under a non-saturated regime. Under a non-saturated regime, both the population correlation and the temporal correlation can detect elevated level of coexpression. Colors represent three different types of external environmental conditions which lead to internal signaling (Steady state, press perturbation, and pulse perturbation). *R*(*τ*) is the cross correlation function with *τ* indicating the time delay. Note that the perturbation is imposed at *t* = 0 in **E** and **H**.

Fig. 4E shows that, under a saturated regime, the population-level correlation between *X* and *Y* can appear even lower under a perturbation, despite the fact that the interaction between *X* and *Y* is activated by an environmental perturbation. On the other hand, under a non-saturated regime, the population-level correlation between the regulator and its target increases (Fig. 4H). It should be noted that under a non-cooperative binding mode (i.e., Hill coefficient equals 1), the population-level correlation will decrease independently under the saturation regime. Therefore, how population-level correlations change relies upon whether a given transcriptional interaction is under a saturated regime or not; population-level correlations can fail to capture such transient environmentally responsive links. These results are robust across parameter configurations (Fig. S8 and Fig. S9). Such a bias can be termed the temporal bias (Yuan et al. 2021). However, the temporal correlation between TF *X* and target gene *Y* is sensitive enough to characterize the environmentally induced activation under both saturated and non-saturated regimes induced by either press or pulse perturbation by the signal, *S*_*x*_ (Fig. 4D and G). These results highlight the likely incidence of false negatives in identifying responsive gene interactions when relying on population-level correlations. While Bar-Joseph and colleagues (Bar-Joseph et al. 2012) have argued that temporal information enable the identification of transient transcription changes, our results suggest that even if transient transcriptional changes of single gene are captured the population correlation analysis can also have low power to identify responsive links because of potential regulatory saturation (as shown in simulation Fig. 4E and empirical data Fig. 3A and 3B), reinforcing the importance of using temporal information to recover environmentally responsive interactions.

However, despite our observation that temporal correlation can robustly detect transient responses of regulatory activity, this approach may obscure the complicated dynamics of regulatory activity over the time course. Specifically, temporal correlations cannot track real-time regulatory activities (Fig. 4D and Fig. 4G). Conversely, population correlation over multiple time-steps may recover the dynamic activity of a regulatory interaction (Fig. 4E and 4H). In the following sections, we will leverage the temporal component of stress response by using both temporal correlations and population correlations over time.

### 3.3 Temporal correlations prioritizing novel candidates in regulating stress responses

We next analysed dynamic transcriptional rewiring through temporal correlation. We examined whether certain TF families affect the activation of regulatory interactions (Fig. S10) and find that, as expected, many TFs with high differential mean MCC in the heat-stress data set are annotated as Heat Shock transcription Factors (HSFs).

Inspecting the relationship between differential gene expression and the differential activity for a given TF here estimated by MCCs of a TF with its target genes reveals that several known HSFs do not independently show strong expression response to the stressor but do show a clear response according to the differential activity. We also find several interesting candidate TFs outside of the HSF family which have high differential regulatory scores but little or no apparent differential expression in pairwise contrasts between control and treatment conditions (Fig. 5A). In the heat data set, the TF OsTCP7 has a differential regulatory score of 0.54 but was not identified as differentially expressed by Wilkins et al. 2016 (Fig. 5C). TCPs are broadly involved in regulating cell proliferation and growth (Martín-Trillo and Cubas 2010) and so OsTCP7 may be an interesting candidate for functional validation in the context of heat stress response.

**Figure 5:**
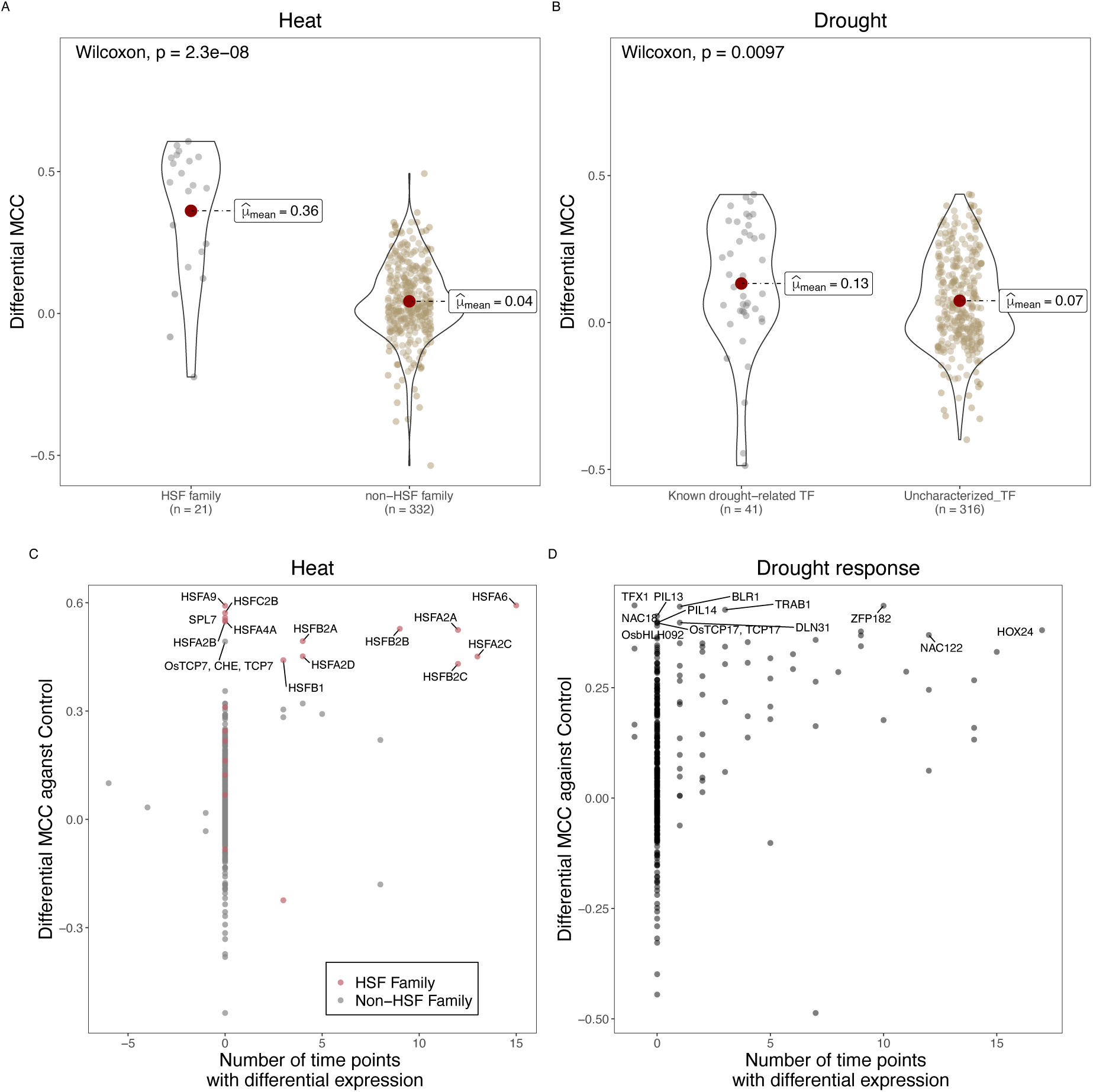
Temporal correlation reveals putative key regulators of stress response. **A**. The average differential Max Correlation Coefficient (MCC) for each regulator in the network prior under heat condition. Violin plots show members of the HSF family TF and non-HSF family TF, respectively. **B**. The average differential MCC for each regulator in the network prior under drought condition. Known drought-related TFs were obtained from https://funricegenes.github.io/, where genes linked with keywords “drought”, “ABA”, and “drought tolerance” were extracted. The average differential MCC is calculated as the averaged MCC changed across conditions. The comparison of heat **C**. and drought **D**. differential expression (the number of time points showing differential expression from the original Wilkins et al. analysis) versus differential MCC. Salmon points denote the Heat Shock Family (HSF) regulators. The number of time points with differential expression is counted for each time point and each genotypes (Maximum number is 4 * 16 = 64). Negative numbers on the x-axis indicate number of time points in which the gene was observed to be downregulated as compared to control conditions.

While the HSF TFs comprise a gene family and are generally interpreted as coordinating plant response to heat stress (von Koskull-Döring et al. 2007), the regulatory control of response to soil drying is more distributed among diverse gene families and regulatory pathways (Joshi et al. 2016, Manna et al. 2020, Des Marais et al. 2012). Our analysis of the drought response data identified several TFs with previously demonstrated roles in rice dehydration response (Fig. 5D). These include HOX24 (Bhattacharjee et al. 2021) and ZFP182 (Huang et al. 2012), both of which were also found by Wilkins et al. to show a strong differential response. Several interesting candidates emerge among the list of TFs which have high differential regulatory score but low differential expression response. One such gene is PIF-Like 12 (Nakamura et al. 2007) which, to our knowledge, has no known role in dehydration response but is paralagous to OsPIL1 which integrates cues from the circadian clock and dehydration signaling to control internode elongation in rice (Todaka et al. 2012). Additional candidate genes with high regulatory scores under elevated temperature or dehydration stress are shown in Table S1 and S2. We hypothesize that the differential activity calculated by temporal correlation could be used to identify novel stress-responsive regulators in this and other systems.

### 3.4 Dynamic TF activity under dehydration conditions reveal signal propagation upon environmental perturbations

Above we showed using stochastic simulations that population correlations may be suitable for estimating the activity of a regulatory link even though they may miss transient interaction changes (Fig. 4). On the other hand, temporal correlation is capable of capturing transient responses of regulatory activities but may miss some important information over the whole time course of treatment since it only gives a single summarized value without possible temporal fluctuation during the time course, leading to a different type of temporal bias. In the rice dataset considered here, temporal correlations do not show a strong signal in detecting drought-responsive TF (Fig. 5B), while temporal correlations do detect heat-responsive TFs (Fig. 5A). To explore the possible reason for this discrepancy, we next analyze TF activities over time under drought treatment by using the population correlation. We find that the population correlation can indeed unveil the dynamic regulatory map in additional layers through temporal correlation.

We first construct a network hierarchy in the TF-only subnetwork from the network prior (Fig. 6A). Since the network has feedback loops (See Supporting Information A.1), we used a generalized bottom-up approach (Yu and Gerstein 2006). In essence, we define all TFs that do not regulate any other TFs as bottom TFs and define the level of the remaining TFs by their shortest distance to a bottom TF. Caveats and the other alternative approach constructing hierarchy are discussed in the Supporting Information (Fig. S12). 89 of the 357 TFs in the network neither regulate other TFs nor are themselves regulated by other TFs; thus, these 89 TFs are not present in the generalized hierarchical structure. The regulatory signal can be amplified and propagated in a top-down manner, which can be observed in the mean expression level and temporal fluctuation (based on the rice transcriptome data) of TFs in the generalized hierarchy, in which TFs in the top layers show lower expression and weaker fluctuations as compared to the bottom TFs (Fig. 6B). Such evidence implies that differential expression analysis may bias towards bottom TFs and other downstream target genes which likely have higher transcript abundances and higher fluctuations that provide the variance necessary to infer signatures of environmental response.

**Figure 6:**
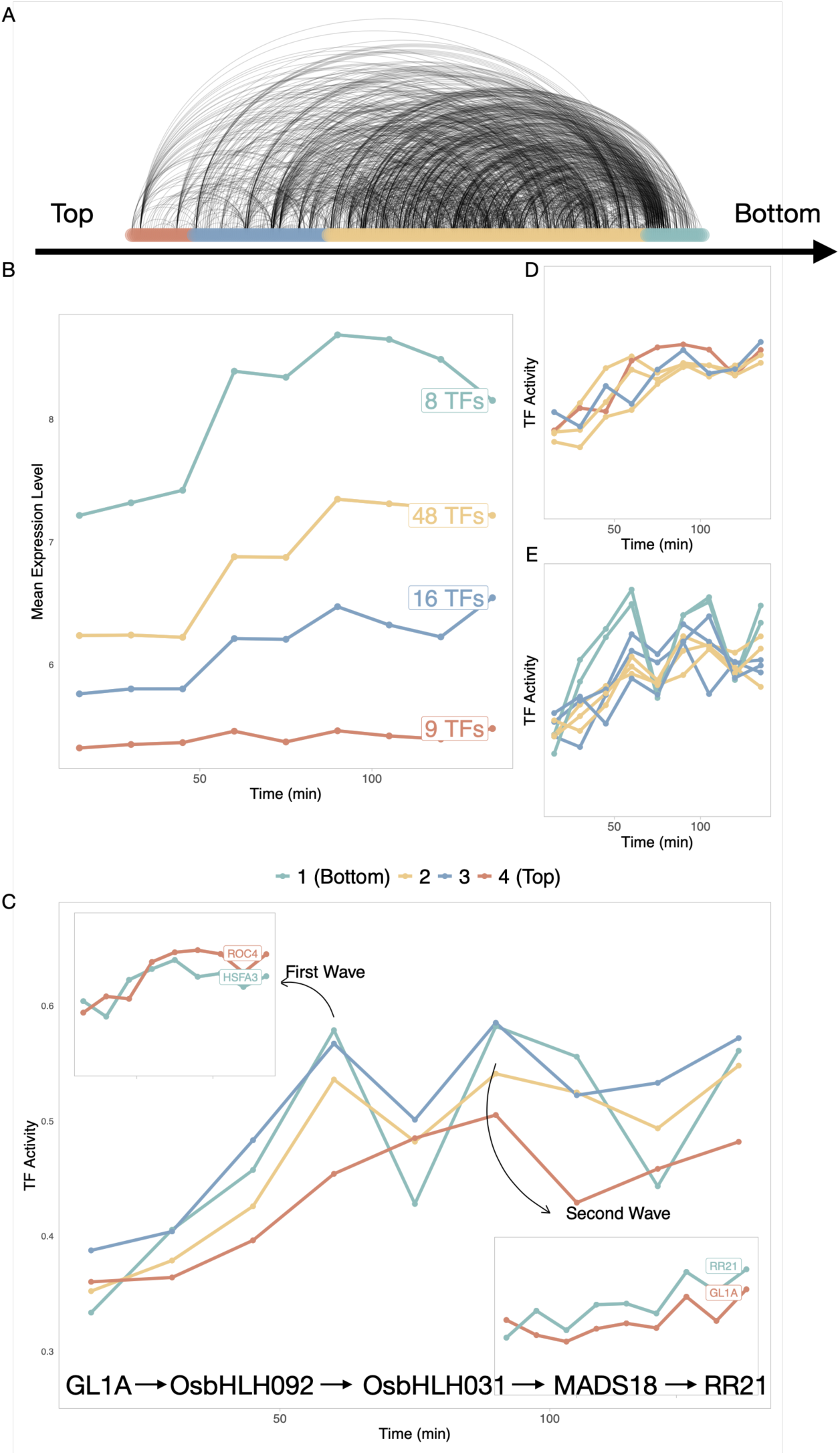
Population correlation over time characterizing the dynamics of Transcription Factor activities under dehydration in a regulatory network hierarchy. **A**. The hierarchical structure of the network prior constructed by a generalized bottom-up approach. Each curve represents a regulatory interaction in the network prior. The color indicates the level of a Transcription Factor (TF) in the hierarchy from top (left) to bottom (right) **B**. Comparison of mean expression value of responsive TFs in the network hierarchy. The label of each line shows the number of TFs within each level. **C**. Dynamic TF activities calculated by average population Pearson Correlation Coefficients of a TF with all its target genes. Two waves of TF activity can be observed; we thus clustered all the TF’s activity within the hierarchy over time with the assistant of STEM (Ernst and Bar-Joseph 2006). Four distinct patterns are shown: 1) Continuously increasing **D**; 2). Driving both two waves (E); 3) and 4) are two groups of TFs drive the first and second regulatory wave separately **Insets**. of **C**.. The shortest path in the **C** shows the regulatory cascade driving the second regulatory wave.

We next examined dynamic TF activities after the drought treatment. A transcription factor’s activity at one time point is calculated as the average population coexpression level (PCC) with all of its target genes at that time point in the static network prior we used above. Therefore, we obtained temporal activities of TFs at control and drought condition. We filtered the TF pools by removing non-responsive TF using paired t test (p < 0.05) comparing temporal activities of a TF across two conditions (although this filtering step has only a moderate effect on the results; See Fig S11). Note that many responsive TFs do not individually exhibit differential expression between control and drought conditions, suggesting that many transcriptional regulations can occur without significantly changing the abundance of the regulators themselves. And, as expected, genes that cannot be detected by differential expression but are identified as responsive TFs by their activities tend to occur in the upper layers of the TF hierarchy. Specifically, 7 out of 9 responsive TFs at top layers do not exhibit differential expression at any time point, while there is only 1 out of 8 responsive TFs showing differential expression. This observation suggests that there is signal amplification through the transcriptional cascade: higher layer TFs and master regulators are more likely to control downstream genes without a detectable change in their own transcript abundances in the data available. Therefore, when analyzing a transcriptome of transcription factors, contrasting regulatory activities (e.g., using coexpression as we did) under stress and control condition may be necessary for prioritizing responsive genes because the signal of differential expression appears to decay along the regulatory hierarchy from top down.

Overall, we observed two regulatory waves for temporal activities (Fig. 6C), which were not prominent in the temporal expression profile considered above (Fig. 6B). We speculate that these two waves may represent distinct phases rice response to the relatively severe drying imposed, perhaps before and after turgor loss occurs (Buckley 2005). We note that Wilkins et al. observed two distinct phases of drying response with respect to carbon assimilation, with a steep decline in assimilation during the initial phase followed by a slower decline beginning around the 60th minute following onset of dehydration stress (Wilkins et al. 2016). We next clustered short time series of single TF activities with STEM (Ernst and Bar-Joseph 2006), which identifies interactions using unsupervised clustering and, in our data, infers the occurence of two regulatory waves along with putative drivers of each wave. We found four distinct groups of TFs (Fig. 6C insets, Fig. 6D and Fig. 6E). Fig. 6D presents a group of TFs with continuously increasing activity over the time course. Fig. 6E, top left inset of Fig. 6C and bottom right inset of Fig. 6C show groups driving both waves, the first wave and the second wave alone respectively. The group of TFs putatively driving the second wave contains two TFs: RR21, from the bottom layer, encodes a putative response regulator involved in cytokinin signaling (Tsai et al. 2012), while GL1A, from the top layer, encodes an ortholog of the Arabidopsis R2R3 MYB transcription factor GL1 (Zheng et al. 2021). A cascade of TFs representing the shortest path between these two TFs were identified in the TF-only network prior (Fig. 6C), showing how regulatory signals are propogated through the network. In summary, instead of temporal correlation, examining population level coexpression afford us opportunity to track the evolving regulatory activity following perturbation.

## 4 DISCUSSION

Many regulatory interactions in TRNs, and thus co-expression relationships inferred from genomescale gene expression measures, are context-dependent (Dunlop et al. 2008, Luscombe et al. 2004). Environmental cues can affect the behavior of regulators by, e.g., changing their abundance or their binding affinity to target DNA sequences, and thereby change their regulatory interactions with other genes and possibly effect whole-organismal response. For instance, an interaction can be inactive simply because the concentration of the regulator is outside its effective range for the target (Dunlop et al. 2008). Notably, even if a regulatory interaction is activated, its regulatory activity can be low as the dose-response curve may be under a saturated regime in which additional units of the regulator do not result in changed activity of its target(s). Alternatively, interactions may be inactive as a result of the chromatin state of target genes, the post-translational modification of the regulator itself, the presence of inhibitory factors, or the absence of co-factors (Toledo and Wahl 2006, Piggot and Hilbert 2004). While these fundamental features of gene regulation are widely appreciated, it is less often considered how such transient processes may confound ecological studies, or how they might be exploited to enhance the interpretative value of the types of transcriptomic datasets now routinely generated by ecologists and evolutionary biologists, which is a central motivation for our study.

Studies in human disease and plant and animal stress response frequently use genome-wide gene expression data to study changes in co-expression changes and network rewiring in response to environmental perturbation (Fukushima 2013, Southworth et al. 2009, de la Fuente 2010, Amar et al. 2013, Choi et al. 2005, Kostka and Spang 2004, Deng et al. 2015, Yan et al. 2019, Cho et al. 2009, Fukushima et al. 2012, de la Fuente 2010, Zeisel et al. 2015, Bhar et al. 2013, Fiannaca et al. 2015). Ultimately, the objective of these studies is to identify the genetic and molecular basis of environmental responses as a means to parameterize models of molecular evolution (Wray et al. 2003), to identify the molecular genetic basis of evolutionary adaptation (Cheviron et al. 2012, Garfield et al. 2013, Carvallo et al. 2011), to understand abnormal regulation in disease states, and to design medical interventions (Southworth et al. 2009, de la Fuente 2010, Amar et al. 2013, Kostka and Spang 2004) and breeding strategies (Fukushima et al. 2012). However, many past studies have relied on population-level statistics to identify differential gene-gene interactions (Cortijo et al. 2020, Fukushima et al. 2012, Deng et al. 2015, Lea et al. 2019). While statistically straightforward, such widely used population-based methods likely miss many dynamic interactions which drive the organismal response, thereby generating an incomplete picture of these complex systems. First, without a static transcriptional network prior, generating a pairwise gene co-expression network and detecting responsive links can lead to false positives as many of links may be indirect and not involve any causal regulatory relationship (Feizi et al. 2013, Barzel and Barabási 2013). Second, population-level statistics are often confounded by individual covariates such as genotype, age, and sex (Parsana et al. 2019, Lea et al. 2019). Third, even within an isogenic homogeneous population, cross sectional population correlations may be confounded by switch-like transitions and ultrasensitivity in gene regulation, thereby failing to characterize the dynamic network rewiring (Fig. 3 and Fig. 4). By contrast, temporal correlation can fail to identify real-time regulatory activity (Fig. 4).

In the present study, we implemented a stochastic simulation of a simple regulatory model under two perturbation regimes. We show that, under a cooperative binding mode (Hill coeffecient *n* > 1, indicating that early-acting TFs enhance bindings of TFs that come later), the population-level co-expression changes of an environmentally induced link depend upon whether the gene regulation is under a saturated regime or not. That is to say, differential population coexpression is neither powerful nor robust in detecting responsive links. Our results also indicate that while population-level correlations may be confounded by saturated regulation, temporal correlations of gene expression time series are robust under both saturated and unsaturated regimes. Hence, while temporal co-expression tends to be coherent upon environmental perturbation, whether the co-expression measured using population statistics becomes coherent or decoherent may depend upon the specific parameters of a given gene pair and the environmental condition. That such potential temporal bias can occur when using cross-sectional data (i.e., population level correlation) has been established in the medical literature (Yuan et al. 2021). Notably, by measuring the activity of 6500 designed promoters using a fluorescence reporter, van Dijk and colleagues (van Dijk et al. 2017) found that the activities of the target promoters can become saturated with increasing abundance of the active form of TFs, and that the pattern becomes more pronounced with more binding sites or higher binding affinity.

Plant responses to environmental stressors are physiologically complex (Bohnert et al. 1995) and often context-dependent and species-specific (Bouzid et al. 2019). Importantly, we found that temporal bias is prominent under heat stress, whereas under drought stress, such bias is not. Furthermore, when removing other covariates (genotypic variation), drought treatments lead to clear patterns of regulatory coherence and increasing phenotypic integration when assessing population correlations, which suggests that regulatory saturation is relatively less common during drought response. We reason that such a distinct pattern reflects the varying etiology of response to different stressors and is largely attributable to the internal environment an organism experienced during stress onset: drying is a fairly gradual process internally, while the heat treatment is a shock – particularly as often implemented in laboratory settings – and a more rapid onset process for an organism. Therefore, we suspect that the drought response is under an unsaturated regime in the rice data studied here and is relatively mild compared to the heat response.

On the contrary, the heat shock treatment is intense and thereby possibly under a saturated regulatory regime in the rice data. Environmental responses that show regulatory saturation may not have been optimized by natural selection: rapid heat shock of the type often imposed in the laboratory seting is likely rare in the wild. Further analysis is needed to examine to what extent this type of saturated regulatory regime is present in real data.

Our observation that co-expression increases with the onset of two stresses in rice is seemingly inconsistent with a recent study which used gene expression data collected from human monocytes to infer population correlation among transcripts (Lea et al. 2019). Several other population-level studies have likewise reported that environmental perturbation may lead to declining co-expression (Southworth et al. 2009, Anglani et al. 2014, Tanner et al. 2022). From a quantitative genetic perspective, a commonly observed result is that phenotypic integration in a population (i.e., the number and strength of significant correlations among traits) increases with environmental stress (Waitt and Levin 1993, Schlichting 1989, Gianoli 2004, García-Verdugo et al. 2009). However, stress-induced decanalization theory (Gibson 2009) suggests that new mutations or stressful environments may disrupt fine-tuned connections in a transcriptional network (Lea et al. 2019). Notably, decanalization has been hypothesized to explain complex traits and human disease (Hu et al. 2016). The degree of stress imposed on the system may dictate whether coherence (or integration) as opposed to decoherence is observed. A possible reconciliation between our results and the decoherence reported by Lea et al, may be that the monocytes studied by Lea et al. experienced a relatively more stressful environment than the rice plants studied by Wilkins et al. Indeed, we recently showed that trait co-variances vary considerably along a single environmental index (Monroe et al. 2021), suggesting that genome-scale regulatory interactions likely transition from coherent to decoherent along such axes. Such gradients of response should be carefully considered during the design of experiments (Poorter et al. 2016).

## 5 Competing Interests

The authors declare that they have no competing interests

## 6 Data Accessibility Statement

The RNA-seq data were retrieved from the Gene Expression Omnibus (www.ncbi.nlm.nih.gov/geo/) under accession number GSE74793.

## Notes

https://www.ncbi.nlm.nih.gov/geo/query/acc.cgi?acc=GSE74793

### Competing Interest Statement

The authors have declared no competing interest.

### Summary of Updates

Fixing figure reference Uploading supporting information

## References

Alon, U., 2019. An introduction to systems biology: design principles of biological circuits. CRC press.

Alonso-Blanco, C., Gomez-Mena, C., Llorente, F., Koornneef, M., Salinas, J., and Martínez-Zapater, J. M. 2005. Genetic and molecular analyses of natural variation indicate cbf2 as a candidate gene for underlying a freezing tolerance quantitative trait locus in arabidopsis. Plant physiology 139:1304–1312.

Alvarez-Fernandez, R., Penfold, C. A., Galvez-Valdivieso, G., Exposito-Rodriguez, M., Stallard, E. J., Bowden, L., Moore, J. D., Mead, A., Davey, P. A., Matthews, J. S., et al. 2020. Time series transcriptomics reveals a bbx32-directed control of dynamic acclimation to high light in mature arabidopsis leaves. bioRxiv.

Amar, D., Safer, H., and Shamir, R. 2013. Dissection of regulatory networks that are altered in disease via differential co-expression. PLoS computational biology 9.

Anglani, R., Creanza, T. M., Liuzzi, V. C., Piepoli, A., Panza, A., Andriulli, A., and Ancona, N. 2014. Loss of connectivity in cancer co-expression networks. PloS one 9.

Bar-Joseph, Z., Gitter, A., and Simon, I. 2012. Studying and modelling dynamic biological processes using time-series gene expression data. Nature Reviews Genetics 13:552–564.

Barzel, B. and Barabási, A.-L. 2013. Network link prediction by global silencing of indirect correlations. Nature biotechnology 31:720–725.

Bechtold, U., Penfold, C. A., Jenkins, D. J., Legaie, R., Moore, J. D., Lawson, T., Matthews, J. S., Vialet-Chabrand, S. R., Baxter, L., Subramaniam, S., et al. 2016. Time-series transcriptomics reveals that agamous-like22 affects primary metabolism and developmental processes in drought-stressed arabidopsis. The Plant Cell 28:345–366.

Bhar, A., Haubrock, M., Mukhopadhyay, A., Maulik, U., Bandyopadhyay, S., and Wingender, E. 2013. Coexpression and coregulation analysis of time-series gene expression data in estrogen-induced breast cancer cell. Algorithms for molecular biology 8:1–11.

Bhattacharjee, A., Srivastava, P. L., Nath, O., and Jain, M. 2021. Genome-wide discovery of oshox24-binding sites and regulation of desiccation stress response in rice. Plant Molecular Biology 105:205–214.

Blüthgen, N., Legewie, S., Herzel, H., and Kholodenko, B., 2007. Mechanisms generating ultrasensitivity, bistability, and oscillations in signal transduction. Pages 282–299 in Introduction to Systems Biology. Springer.

Bohnert, H. J., Nelson, D. E., and Jensen, R. G. 1995. Adaptations to environmental stresses. The plant cell 7:1099.

Bouzid, M., He, F., Schmitz, G., Häusler, R., Weber, A., Mettler-Altmann, T., and De Meaux, J. 2019. Arabidopsis species deploy distinct strategies to cope with drought stress. Annals of botany 124:27–40.

Bradshaw, A. D. 1965. Evolutionary significance of phenotypic plasticity in plants. Advances in genetics 13:115–155.

Briat, C., Gupta, A., and Khammash, M. 2016. Antithetic integral feedback ensures robust perfect adaptation in noisy biomolecular networks. Cell systems 2:15–26.

Buchanan, M., Caldarelli, G., De Los Rios, P., Rao, F., and Vendruscolo, M., 2010. Networks in cell biology. Cambridge University Press.

Buckley, T. N. 2005. The control of stomata by water balance. New phytologist 168:275–292.

Carroll, S. B. 2008. Evo-devo and an expanding evolutionary synthesis: a genetic theory of morphological evolution. Cell 134:25–36.

Carvallo, M. A., Pino, M.-T., Jeknić, Z., Zou, C., Doherty, C. J., Shiu, S.-H., Chen, T. H., and Thomashow, M. F. 2011. A comparison of the low temperature transcriptomes and cbf regulons of three plant species that differ in freezing tolerance: Solanum commersonii, solanum tuberosum, and arabidopsis thaliana. Journal of experimental botany 62:3807–3819.

Chapin, F. S. 1991. Integrated responses of plants to stress. BioScience 41:29–36.

Cheviron, Z. A., Bachman, G. C., Connaty, A. D., McClelland, G. B., and Storz, J. F. 2012. Regulatory changes contribute to the adaptive enhancement of thermogenic capacity in high-altitude deer mice. Proceedings of the national academy of sciences 109:8635–8640.

Cho, S. B., Kim, J., and Kim, J. H. 2009. Identifying set-wise differential co-expression in gene expression microarray data. BMC bioinformatics 10:109.

Choi, J. K., Yu, U., Yoo, O. J., and Kim, S. 2005. Differential coexpression analysis using microarray data and its application to human cancer. Bioinformatics 21:4348–4355.

Chu, D., Zabet, N. R., and Mitavskiy, B. 2009. Models of transcription factor binding: sensitivity of activation functions to model assumptions. Journal of Theoretical Biology 257:419–429.

Cortijo, S., Bhattarai, M., Locke, J. C., and Ahnert, S. E. 2020. Co-expression networks from gene expression variability between genetically identical seedlings can reveal novel regulatory relationships. Frontiers in plant science 11.

de la Fuente, A. 2010. From ‘differential expression’to ‘differential networking’–identification of dysfunctional regulatory networks in diseases. Trends in genetics 26:326–333.

Deng, S.-P., Zhu, L., and Huang, D.-S., 2015. Mining the bladder cancer-associated genes by an integrated strategy for the construction and analysis of differential co-expression networks. in BMC genomics, volume 16, page S4. BioMed Central.

Des Marais, D. L., Hernandez, K. M., and Juenger, T. E. 2013. Genotype-by-environment interaction and plasticity: exploring genomic responses of plants to the abiotic environment. Annual Review of Ecology, Evolution, and Systematics 44:5–29.

Des Marais, D. L., McKay, J. K., Richards, J. H., Sen, S., Wayne, T., and Juenger, T. E. 2012. Physiological genomics of response to soil drying in diverse arabidopsis accessions. The Plant Cell 24:893–914.

Dunlop, M. J., Cox, R. S., Levine, J. H., Murray, R. M., and Elowitz, M. B. 2008. Regulatory activity revealed by dynamic correlations in gene expression noise. Nature genetics 40:1493–1498.

Ernst, J. and Bar-Joseph, Z. 2006. Stem: a tool for the analysis of short time series gene expression data. BMC bioinformatics 7:1–11.

Feizi, S., Marbach, D., Médard, M., and Kellis, M. 2013. Network deconvolution as a general method to distinguish direct dependencies in networks. Nature biotechnology 31:726–733.

Ferrari, C., Proost, S., Ruprecht, C., and Mutwil, M. 2018. Phytonet: comparative co-expression network analyses across phytoplankton and land plants. Nucleic acids research 46:W76–W83.

Fiannaca, A., La Rosa, M., La Paglia, L., Rizzo, R., and Urso, A. 2015. Analysis of mirna expression profiles in breast cancer using biclustering. BMC bioinformatics 16:1–11.

Fu, X., Sun, Y., Wang, J., Xing, Q., Zou, J., Li, R., Wang, Z., Wang, S., Hu, X., Zhang, L., et al. 2014. Sequencing-based gene network analysis provides a core set of gene resource for understanding thermal adaptation in z hikong scallop c hlamys farreri. Molecular Ecology Resources 14:184–198.

Fukao, T., Yeung, E., and Bailey-Serres, J. 2011. The submergence tolerance regulator sub1a mediates crosstalk between submergence and drought tolerance in rice. The Plant Cell 23:412–427.

Fukushima, A. 2013. Diffcorr: an r package to analyze and visualize differential correlations in biological networks. Gene 518:209–214.

Fukushima, A., Nishizawa, T., Hayakumo, M., Hikosaka, S., Saito, K., Goto, E., and Kusano, M. 2012. Exploring tomato gene functions based on coexpression modules using graph clustering and differential coexpression approaches. Plant physiology 158:1487–1502.

Gao, Q., Ho, C., Jia, Y., Li, J. J., and Huang, H. 2012. Bi c lustering of li near p atterns in gene expression data. Journal of Computational Biology 19:619–631.

García-Verdugo, C., Granado-Yela, C., Manrique, E., Rubio de Casas, R., and Balaguer, L. 2009. Phenotypic plasticity and integration across the canopy of olea europaea subsp. guanchica (oleaceae) in populations with different wind exposures. American Journal of Botany 96:1454–1461.

Garfield, D. A., Runcie, D. E., Babbitt, C. C., Haygood, R., Nielsen, W. J., and Wray, G. A. 2013. The impact of gene expression variation on the robustness and evolvability of a developmental gene regulatory network. PLoS biology 11:e1001696.

Gargouri, M., Park, J.-J., Holguin, F. O., Kim, M.-J., Wang, H., Deshpande, R. R., Shachar-Hill, Y., Hicks, L. M., and Gang, D. R. 2015. Identification of regulatory network hubs that control lipid metabolism in chlamydomonas reinhardtii. Journal of experimental botany 66:4551–4566.

Gerstein, M. B., Kundaje, A., Hariharan, M., Landt, S. G., Yan, K.-K., Cheng, C., Mu, X. J., Khurana, E., Rozowsky, J., Alexander, R., et al. 2012. Architecture of the human regulatory network derived from encode data. Nature 489:91–100.

Gianoli, E. 2004. Plasticity of traits and correlations in two populations of convolvulus arvensis (convolvulaceae) differing in environmental heterogeneity. International Journal of Plant Sciences 165:825–832.

Gianoli, E. and Palacio-López, K. 2009. Phenotypic integration may constrain phenotypic plasticity in plants. Oikos 118:1924–1928.

Gibert, P., Debat, V., and Ghalambor, C. K. 2019. Phenotypic plasticity, global change, and the speed of adaptive evolution. Current opinion in insect science 35:34–40.

Gibson, G. 2008. The environmental contribution to gene expression profiles. Nature reviews genetics 9:575–581.

Gibson, G. 2009. Decanalization and the origin of complex disease. Nature Reviews Genetics 10:134–140.

Gillespie, D. T. 1977. Exact stochastic simulation of coupled chemical reactions. The journal of physical chemistry 81:2340–2361.

Greenham, K., Guadagno, C. R., Gehan, M. A., Mockler, T. C., Weinig, C., Ewers, B. E., and McClung, C. R. 2017. Temporal network analysis identifies early physiological and transcriptomic indicators of mild drought in brassica rapa. Elife 6:e29655.

Hirata, H., Yoshiura, S., Ohtsuka, T., Bessho, Y., Harada, T., Yoshikawa, K., and Kageyama, R. 2002. Oscillatory expression of the bhlh factor hes1 regulated by a negative feedback loop. Science 298:840–843.

Horn, P. J., Liu, J., Cocuron, J.-C., McGlew, K., Thrower, N. A., Larson, M., Lu, C., Alonso, A. P., and Ohlrogge, J. 2016. Identification of multiple lipid genes with modifications in expression and sequence associated with the evolution of hydroxy fatty acid accumulation in physaria fendleri. The Plant Journal 86:322–348.

Hu, J. X., Thomas, C. E., and Brunak, S. 2016. Network biology concepts in complex disease comorbidities. Nature Reviews Genetics 17:615.

Huang, J., Sun, S., Xu, D., Lan, H., Sun, H., Wang, Z., Bao, Y., Wang, J., Tang, H., and Zhang, H. 2012. A tfiiia-type zinc finger protein confers multiple abiotic stress tolerances in transgenic rice (oryza sativa l.). Plant molecular biology 80:337–350.

Joshi, R., Wani, S. H., Singh, B., Bohra, A., Dar, Z. A., Lone, A. A., Pareek, A., and Singla-Pareek, S. L. 2016. Transcription factors and plants response to drought stress: current understanding and future directions. Frontiers in Plant Science 7:1029.

Kostka, D. and Spang, R. 2004. Finding disease specific alterations in the co-expression of genes. Bioinformatics 20:i194–i199.

Langfelder, P. and Horvath, S. 2008. Wgcna: an r package for weighted correlation network analysis. BMC bioinformatics 9:559.

Lea, A., Subramaniam, M., Ko, A., Lehtimäki, T., Raitoharju, E., Kähönen, M., Seppälä, I., Mononen, N., Raitakari, O. T., Ala-Korpela, M., et al. 2019. Genetic and environmental perturbations lead to regulatory decoherence. eLife 8:e40538.

Lewis, J. 2003. Autoinhibition with transcriptional delay: a simple mechanism for the zebrafish somitogenesis oscillator. Current Biology 13:1398–1408.

Love, M. I., Huber, W., and Anders, S. 2014. Moderated estimation of fold change and dispersion for rna-seq data with deseq2. Genome biology 15:1–21.

Luscombe, N. M., Babu, M. M., Yu, H., Snyder, M., Teichmann, S. A., and Gerstein, M. 2004. Genomic analysis of regulatory network dynamics reveals large topological changes. Nature 431:308–312.

Ma, W., Trusina, A., El-Samad, H., Lim, W. A., and Tang, C. 2009. Defining network topologies that can achieve biochemical adaptation. Cell 138:760–773.

Manna, M., Thakur, T., Chirom, O., Mandlik, R., Deshmukh, R., and Salvi, P. 2020. Transcription factors as key molecular target to strengthen the drought stress tolerance in plants. Physiologia Plantarum.

Martín-Trillo, M. and Cubas, P. 2010. Tcp genes: a family snapshot ten years later. Trends in plant science 15:31–39.

Monaco, G., van Dam, S., Ribeiro, J. L. C. N., Larbi, A., and de Magalhães, J. P. 2015. A comparison of human and mouse gene co-expression networks reveals conservation and divergence at the tissue, pathway and disease levels. BMC evolutionary biology 15:1–14.

Monroe, J., Cai, H., and Des Marais, D. L. 2021. Diversity in non-linear responses to soil moisture shapes evolutionary constraints in brachypodium. G3 Genes| Genomes| Genetics.

Nakamura, Y., Kato, T., Yamashino, T., Murakami, M., and Mizuno, T. 2007. Characterization of a set of phytochrome-interacting factor-like bhlh proteins in oryza sativa. Bioscience, Biotechnology, and Biochemistry 71:1183–1191.

Nicotra, A. B., Atkin, O. K., Bonser, S. P., Davidson, A. M., Finnegan, E. J., Mathesius, U., Poot, P., Purugganan, M. D., Richards, C. L., Valladares, F., et al. 2010. Plant phenotypic plasticity in a changing climate. Trends in plant science 15:684–692.

Ohama, N., Sato, H., Shinozaki, K., and Yamaguchi-Shinozaki, K. 2017. Transcriptional regulatory network of plant heat stress response. Trends in plant science 22:53–65.

Oostra, V., Saastamoinen, M., Zwaan, B. J., and Wheat, C. W. 2018. Strong phenotypic plasticity limits potential for evolutionary responses to climate change. Nature communications 9:1–11.

Palakurty, S. X., Stinchcombe, J. R., and Afkhami, M. E. 2018. Cooperation and coexpression: how coexpression networks shift in response to multiple mutualists. Molecular ecology 27:1860–1873.

Parsana, P., Ruberman, C., Jaffe, A. E., Schatz, M. C., Battle, A., and Leek, J. T. 2019. Addressing confounding artifacts in reconstruction of gene co-expression networks. Genome biology 20:1–6.

Piggot, P. J. and Hilbert, D. W. 2004. Sporulation of bacillus subtilis. Current opinion in microbiology 7:579–586.

Pigliucci, M. 2003. Phenotypic integration: studying the ecology and evolution of complex phenotypes. Ecology letters 6:265–272.

Pigliucci, M. and Preston, K., 2004. Phenotypic integration: studying the ecology and evolution of complex phenotypes. Oxford University Press.

Poorter, H., Fiorani, F., Pieruschka, R., Wojciechowski, T., van der Putten, W. H., Kleyer, M., Schurr, U., and Postma, J. 2016. Pampered inside, pestered outside? differences and similarities between plants growing in controlled conditions and in the field. New Phytologist 212:838–855.

Riaño-Pachón, D. M., Ruzicic, S., Dreyer, I., and Mueller-Roeber, B. 2007. Plntfdb: an integrative plant transcription factor database. BMC bioinformatics 8:1–10.

Ruprecht, C., Proost, S., Hernandez-Coronado, M., Ortiz-Ramirez, C., Lang, D., Rensing, S. A., Becker, J. D., Vandepoele, K., and Mutwil, M. 2017a. Phylogenomic analysis of gene co-expression networks reveals the evolution of functional modules. The Plant Journal 90:447–465.

Ruprecht, C., Vaid, N., Proost, S., Persson, S., and Mutwil, M. 2017b. Beyond genomics: studying evolution with gene coexpression networks. Trends in Plant Science 22:298–307.

Schlichting, C. D. 1989. Phenotypic integration and environmental change. BioScience 39:460.

Schneider, R. F., Li, Y., Meyer, A., and Gunter, H. M. 2014. Regulatory gene networks that shape the development of adaptive phenotypic plasticity in a cichlid fish. Molecular ecology 23:4511–4526.

Smith, H. 1990. Signal perception, differential expression within multigene families and the molecular basis of phenotypic plasticity. Plant, Cell & Environment 13:585–594.

Song, L., Huang, S.-s. C., Wise, A., Castanon, R., Nery, J. R., Chen, H., Watanabe, M., Thomas, J., Bar-Joseph, Z., and Ecker, J. R. 2016. A transcription factor hierarchy defines an environmental stress response network. Science 354:aag1550.

Southworth, L. K., Owen, A. B., and Kim, S. K. 2009. Aging mice show a decreasing correlation of gene expression within genetic modules. PLoS genetics 5.

Stotz, G. C., Salgado-Luarte, C., Escobedo, V. M., Valladares, F., and Gianoli, E. 2021. Global trends in phenotypic plasticity of plants. Ecology Letters 24:2267–2281.

Tanner, R. L., Gleason, L. U., and Dowd, W. W. 2022. Environment-driven shifts in inter-individual variation and phenotypic integration within subnetworks of the mussel transcriptome and proteome. Molecular ecology.

Todaka, D., Nakashima, K., Maruyama, K., Kidokoro, S., Osakabe, Y., Ito, Y., Matsukura, S., Fujita, Y., Yoshiwara, K., Ohme-Takagi, M., et al. 2012. Rice phytochrome-interacting factor-like protein ospil1 functions as a key regulator of internode elongation and induces a morphological response to drought stress. Proceedings of the National Academy of Sciences 109:15947–15952.

Toledo, F. and Wahl, G. M. 2006. Regulating the p53 pathway: in vitro hypotheses, in vivo veritas. Nature Reviews Cancer 6:909–923.

Tsai, Y.-C., Weir, N. R., Hill, K., Zhang, W., Kim, H. J., Shiu, S.-H., Schaller, G. E., and Kieber, J. J. 2012. Characterization of genes involved in cytokinin signaling and metabolism from rice. Plant Physiology 158:1666–1684.

van Dijk, D., Sharon, E., Lotan-Pompan, M., Weinberger, A., Segal, E., and Carey, L. B. 2017. Large-scale mapping of gene regulatory logic reveals context-dependent repression by transcriptional activators. Genome research 27:87–94.

Varala, K., Marshall-Colón, A., Cirrone, J., Brooks, M. D., Pasquino, A. V., Léran, S., Mittal, S., Rock, T. M., Edwards, M. B., Kim, G. J., et al. 2018. Temporal transcriptional logic of dynamic regulatory networks underlying nitrogen signaling and use in plants. Proceedings of the National Academy of Sciences 115:6494–6499.

Villamil, C. I. 2018. Phenotypic integration of the cervical vertebrae in the hominoidea (primates). Evolution 72:490–517.

von Koskull-Döring, P., Scharf, K.-D., and Nover, L. 2007. The diversity of plant heat stress transcription factors. Trends in plant science 12:452–457.

Waitt, D. E. and Levin, D. A. 1993. Phenotypic integration and plastic correlations in phlox drummondii (polemoniaceae). American Journal of Botany 80:1224–1233.

Wang, J., Wu, G., Chen, L., and Zhang, W. 2013. Cross-species transcriptional network analysis reveals conservation and variation in response to metal stress in cyanobacteria. BMC genomics 14:1–11.

Wang, W., Vinocur, B., Shoseyov, O., and Altman, A. 2004. Role of plant heat-shock proteins and molecular chaperones in the abiotic stress response. Trends in plant science 9:244–252.

Weirauch, M. T., Yang, A., Albu, M., Cote, A. G., Montenegro-Montero, A., Drewe, P., Najafabadi, H. S., Lambert, S. A., Mann, I., Cook, K., et al. 2014. Determination and inference of eukaryotic transcription factor sequence specificity. Cell 158:1431–1443.

West-Eberhard, M. J., 2003. Developmental plasticity and evolution. Oxford University Press.

Wilkins, O., Hafemeister, C., Plessis, A., Holloway-Phillips, M.-M., Pham, G. M., Nicotra, A. B., Gregorio, G. B., Jagadish, S. K., Septiningsih, E. M., Bonneau, R., et al. 2016. Egrins (environmental gene regulatory influence networks) in rice that function in the response to water deficit, high temperature, and agricultural environments. The Plant Cell 28:2365–2384.

Windram, O., Madhou, P., McHattie, S., Hill, C., Hickman, R., Cooke, E., Jenkins, D. J., Penfold, C. A., Baxter, L., Breeze, E., et al. 2012. Arabidopsis defense against botrytis cinerea: chronology and regulation deciphered by high-resolution temporal transcriptomic analysis. The Plant Cell 24:3530–3557.

Wray, G. A., Hahn, M. W., Abouheif, E., Balhoff, J. P., Pizer, M., Rockman, M. V., and Romano, L. A. 2003. The evolution of transcriptional regulation in eukaryotes. Molecular biology and evolution 20:1377–1419.

Yan, Q., Wu, F., Yan, Z., Li, J., Ma, T., Zhang, Y., Zhao, Y., Wang, Y., and Zhang, J. 2019. Differential co-expression networks of long non-coding rnas and mrnas in cleistogenes songorica under water stress and during recovery. BMC plant biology 19:1–19.

Yano, M., Katayose, Y., Ashikari, M., Yamanouchi, U., Monna, L., Fuse, T., Baba, T., Yamamoto, K., Umehara, Y., Nagamura, Y., et al. 2000. Hd1, a major photoperiod sensitivity quantitative trait locus in rice, is closely related to the arabidopsis flowering time gene constans. The Plant Cell 12:2473–2483.

Yeung, E., van Veen, H., Vashisht, D., Paiva, A. L. S., Hummel, M., Rankenberg, T., Steffens, B., Steffen-Heins, A., Sauter, M., de Vries, M., et al. 2018. A stress recovery signaling network for enhanced flooding tolerance in arabidopsis thaliana. Proceedings of the National Academy of Sciences 115:E6085–E6094.

Yu, H. and Gerstein, M. 2006. Genomic analysis of the hierarchical structure of regulatory networks. Proceedings of the National Academy of Sciences 103:14724–14731.

Yuan, W., Beaulieu-Jones, B. K., Yu, K.-H., Lipnick, S. L., Palmer, N., Loscalzo, J., Cai, T., and Kohane, sI. S. 2021. Temporal bias in case-control design: preventing reliable predictions of the future. Nature Communications 12:1107. URL https://doi.org/10.1038/s41467-021-21390-2.

Zander, M., Lewsey, M. G., Clark, N. M., Yin, L., Bartlett, A., Guzmán, J. P. S., Hann, E., Langford, A. E., Jow, B., Wise, A., et al. 2020. Integrated multi-omics framework of the plant response to jasmonic acid. Nature Plants 6:290–302.

Zeisel, A., Muñoz-Manchado, A. B., Codeluppi, S., Lönnerberg, P., La Manno, G., Juréus, A., Marques, S., Munguba, H., He, L., Betsholtz, C., et al. 2015. Cell types in the mouse cortex and hippocampus revealed by single-cell rna-seq. Science 347:1138–1142.

Zhao, X., Yu, H., Kong, L., and Li, Q. 2016. Gene co-expression network analysis reveals the correlation patterns among genes in euryhaline adaptation of crassostrea gigas. Marine Biotechnology 18:535–544.

Zheng, K., Wang, X., Wang, Y., and Wang, S. 2021. Conserved and non-conserved functions of the rice homologs of the arabidopsis trichome initiation-regulating mbw complex proteins. BMC plant biology 21:1–15.

