## Supporting information for "Revisiting regulatory decoherence and phenotypic integration: accounting for temporal bias in co-expression analyses"

### A Supporting Information

#### A.1 Network topology and construction of TF hierarchies

The static network prior comprised of 38137 edges, with 1611 edges and 357 TFs in the TF-only subnetwork. The feedback loop number contained in the network are 7 (2-node loop), 19 (three nodes), 39 (four nodes), and 82 (five nodes). Thus, the static network is not a tree-based network. To construct the hierarchy for TF, we utilized two generalized hierarchy approaches. 1). Bottom-up Breadth-First Search, where a TF is at the bottom level if and only if it does not regulate other TFs. Starting from each bottom TF, a non-bottom TF's hierarchy is defined as its shortest distance from any bottom one (Yu and Gerstein 2006). 2). Height-based approach using out- and in-degree of a TF. Hierarchy height of TFs was calculated as previously described (Gerstein et al. 2012):

$$h = \frac{O - I}{O + I} \tag{S1}$$

where $O$ and $I$ are out-degree and in-degree of a TF. We here categorized the height into four levels:

$$\begin{cases} 4, & h = 1 \\ 3, & 0 < h < 1 \\ 2, & -1 < h \leq 0 \\ 1, & h = -1 \end{cases} \tag{S2}$$

Since the network contains feedback loops and feedforward motifs, it is difficult to find the best way to reconstruct the hierarchy. Notably, for instance, there exist two odd nodes in the top layer through Bottom-up Breadth-first search. These two nodes have only two out-degree in the TF-only subnetwork but are regulated by 24 and 27 other TFs respectively. In fact, these two approaches have similar global trends (Fig. S12).

#### A.2 Max lag time specification

The expression relationship observed between genes in a time series sample may be caused by the time lag inherent in molecular interactions, in this case, transcriptional regulation. Such time lag reflects the time required for a TF's activity to influence the expression of its target genes because transcription and translation take place over non-negligible time periods.

The time delay that possesses the max correlation is defined as $\tau_{reg}$, representing the approximate time delay that occurs between a given regulatory-target pair. The max delay is set as 1, which in the current dataset represents a 15 min time interval. We chose a 15 minute interval as the max delay period because several studies have reported that the time duration of transcription and translation is within 30 min (Hirata et al. 2002, Lewis 2003). We also conducted robust tests with delay time 15 and 30 min. The results are qualitatively similar in terms of the regulatory coherence and downstream analysis (Fig. S3).

#### B Supplement Tables

**Table S1: Gene ontology and information of candidate responsive regulators showing high differential regulatory scores following drought perturbations**

**Table S2: Gene ontology and information of candidate responsive regulators showing high differential regulatory scores following heat perturbations**

### C Supplement Figures

#### C.1 Supplement figures for main Figure 2

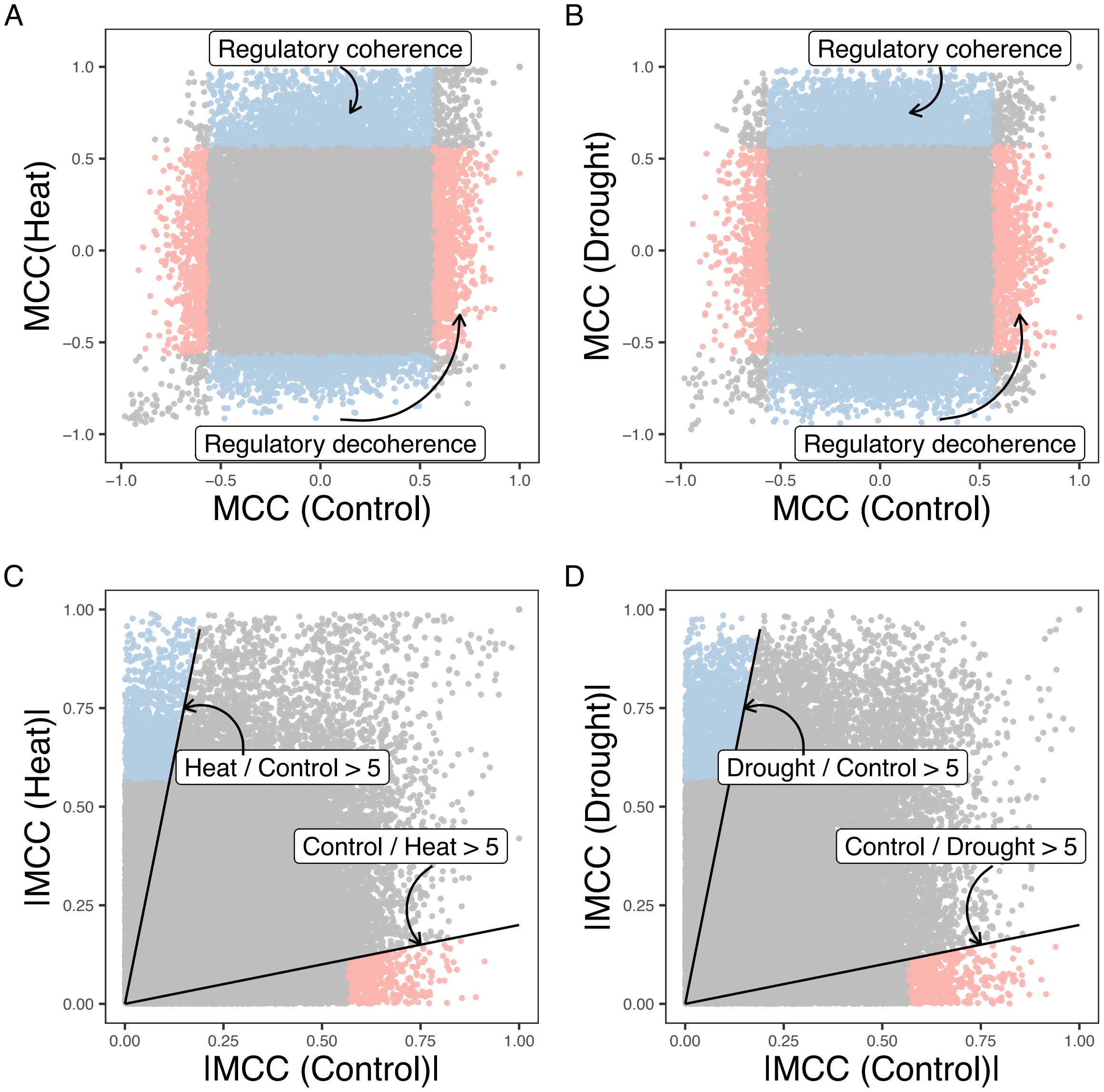


Figure S1: **Environmental perturbations lead to more regulatory coherence. A.** Comparison of the Max Cross Correlation (MCC) for each regulator-target pair under control condition against heat condition. **B.** Comparisons of MCC for each regulator-target pair under control condition against soil drying condition. The regulator-target pairs that are not significant in each condition are in grey, for which the MCC cutoff is 0.567 (corresponding to a p-value of 0.05). The red and blue points highlight the pairs that showing regulatory decoherence and regulatory coherence, respectively. **C.** and **D.** show the absolute value of each regulator-target pair with solid lines indicating that the ratio between regulatory scores under control and perturbed conditions is larger than 5.

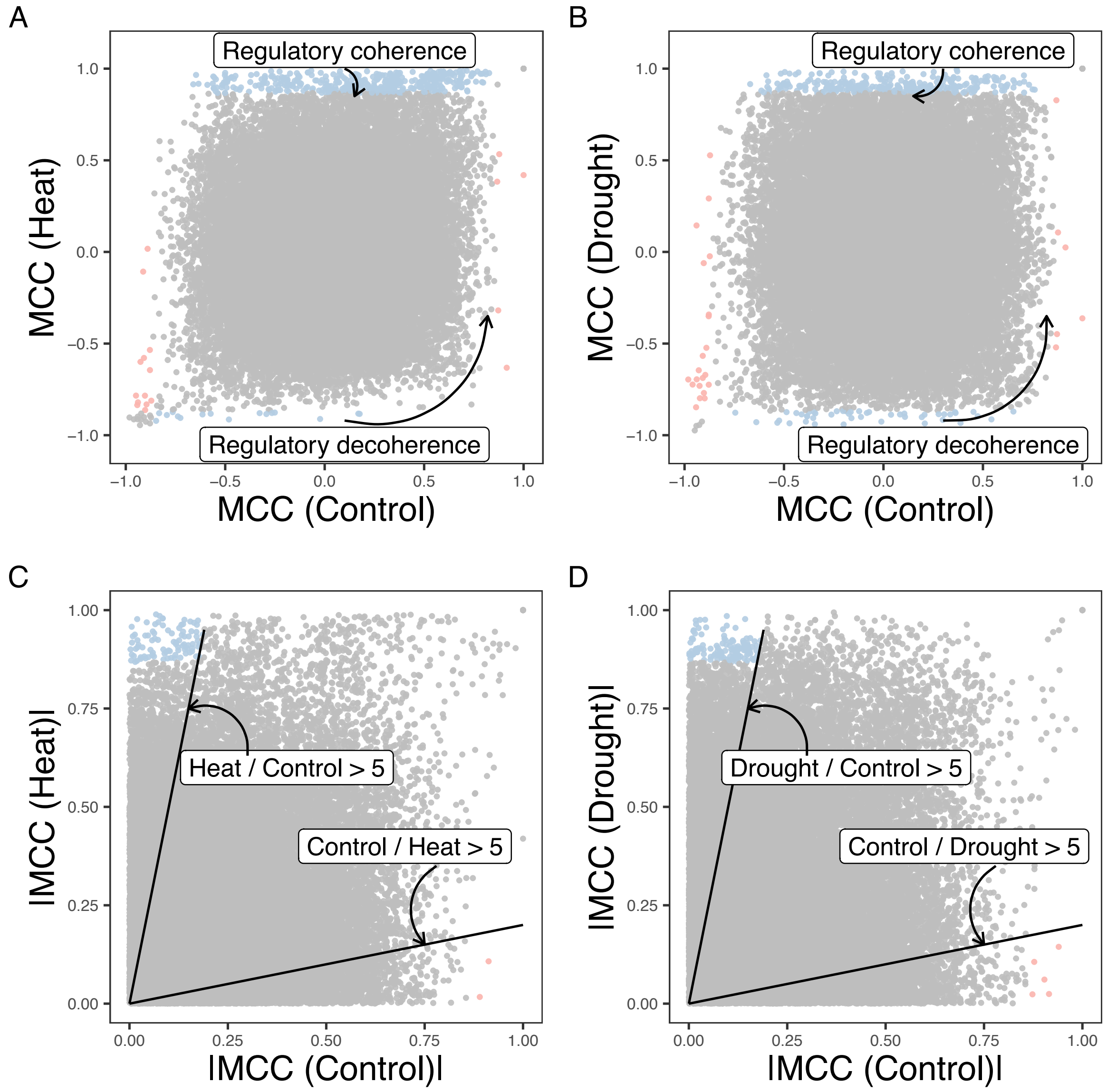


Figure S2: **Environmental perturbations lead to more regulatory coherence. A.** Comparison of the Max Cross Correlation (MCC) for each regulator-target pair under control condition against heat condition. **B.** Comparisons of MCC for each regulator-target pair under control condition against soil drying condition. The regulator-target pairs that are not significant in each condition are in grey, for which the cutoff is 0.867 (corresponding to a p-value of 0.001). The red and blue label highlight the pairs that show regulatory decoherence and regulatory coherence, respectively. **C.** and **D.** show the absolute value of each regulator-target pair with solid lines indicating that the ratio between regulatory scores under control and perturbed conditions is larger than 5.

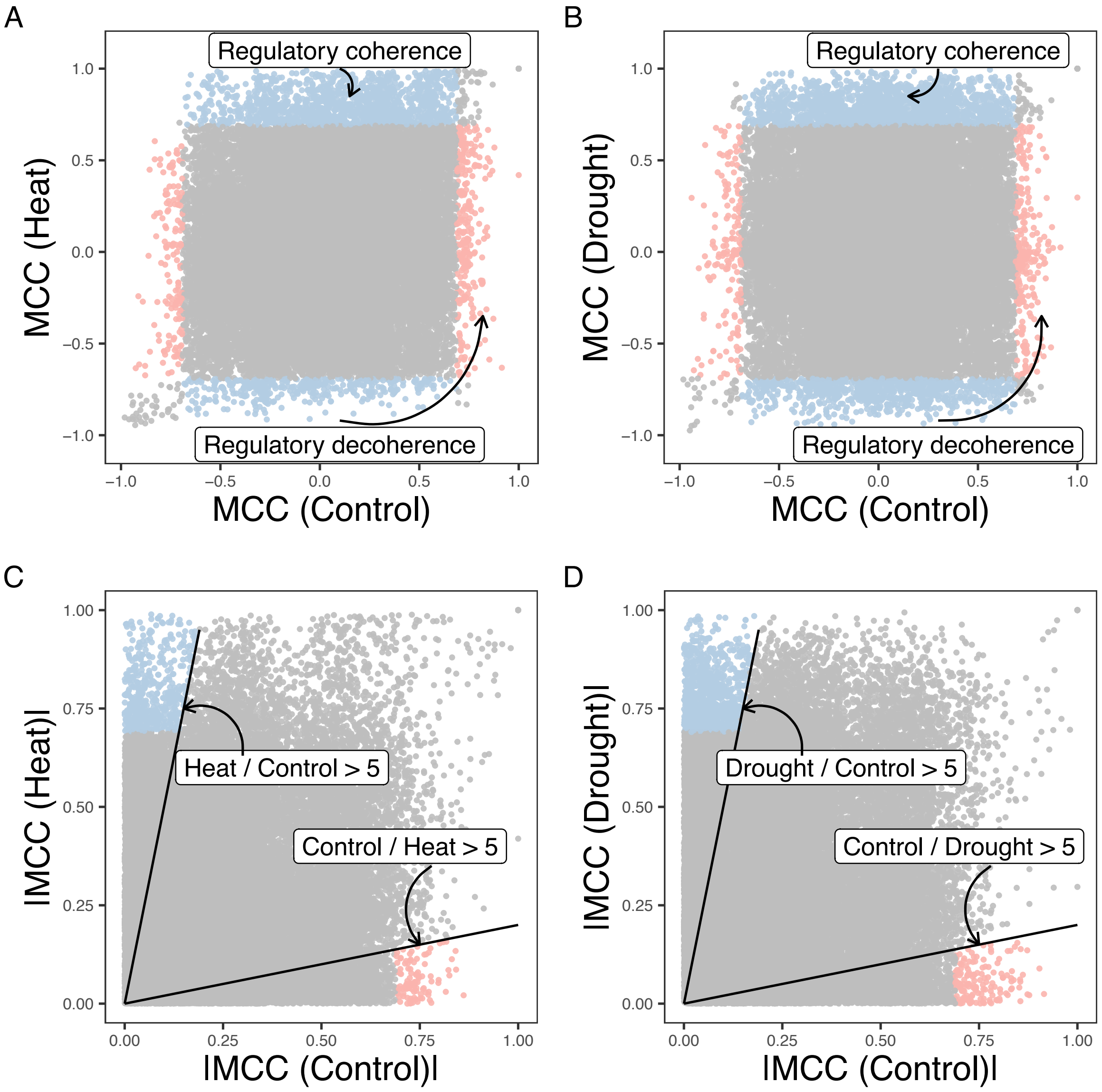


Figure S3: **Environmental perturbations lead to more regulatory coherence.** **A.** Comparison of the Max Cross Correlation (MCC) with max lag time = 2 for each regulator-target pair under control condition against heat condition. **B.** Comparisons of MCC max lag time = 2 for each regulator-target pair under control condition against soil drying condition. The regulator-target pairs that are not significant on both conditions are in grey, for which the cutoff is 0.69 (corresponding to a p-value of 0.01). The red and blue label highlight the pairs that showing regulatory decoherence and regulatory coherence respectively. **C.** and **D.** show the absolute value of each regulator-target pair with solid lines indicating that the ratio between regulatory scores under control and perturbed conditions is larger than 5.

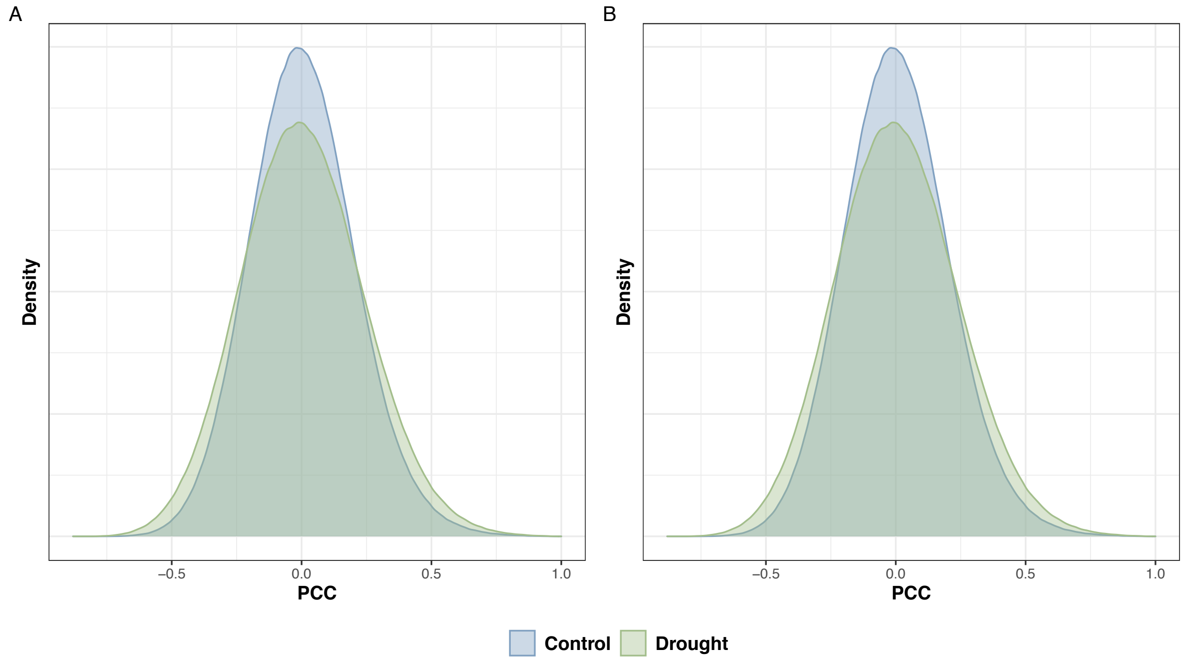


Figure S4: **Drought responses show less temporal bias in *Brachypodium* dataset** Pearson Correlation Coefficient (PCC) distribution for pairwise gene pairs. **A** and **B** are two genotypes (BD21 and BD3-1) respectively. Each genotype contains 16 biological replicates under each condition (drought and control). We ask whether patterns of co-expression differed between leaves assayed at control versus drought treatment. We limit our gene pools by only focusing on differential expression genes (FDR = 0.1 %). Differential expression genes are identified through DESeq2. (Love et al. 2014)

#### C.2 Supplement figures for main Figure 3

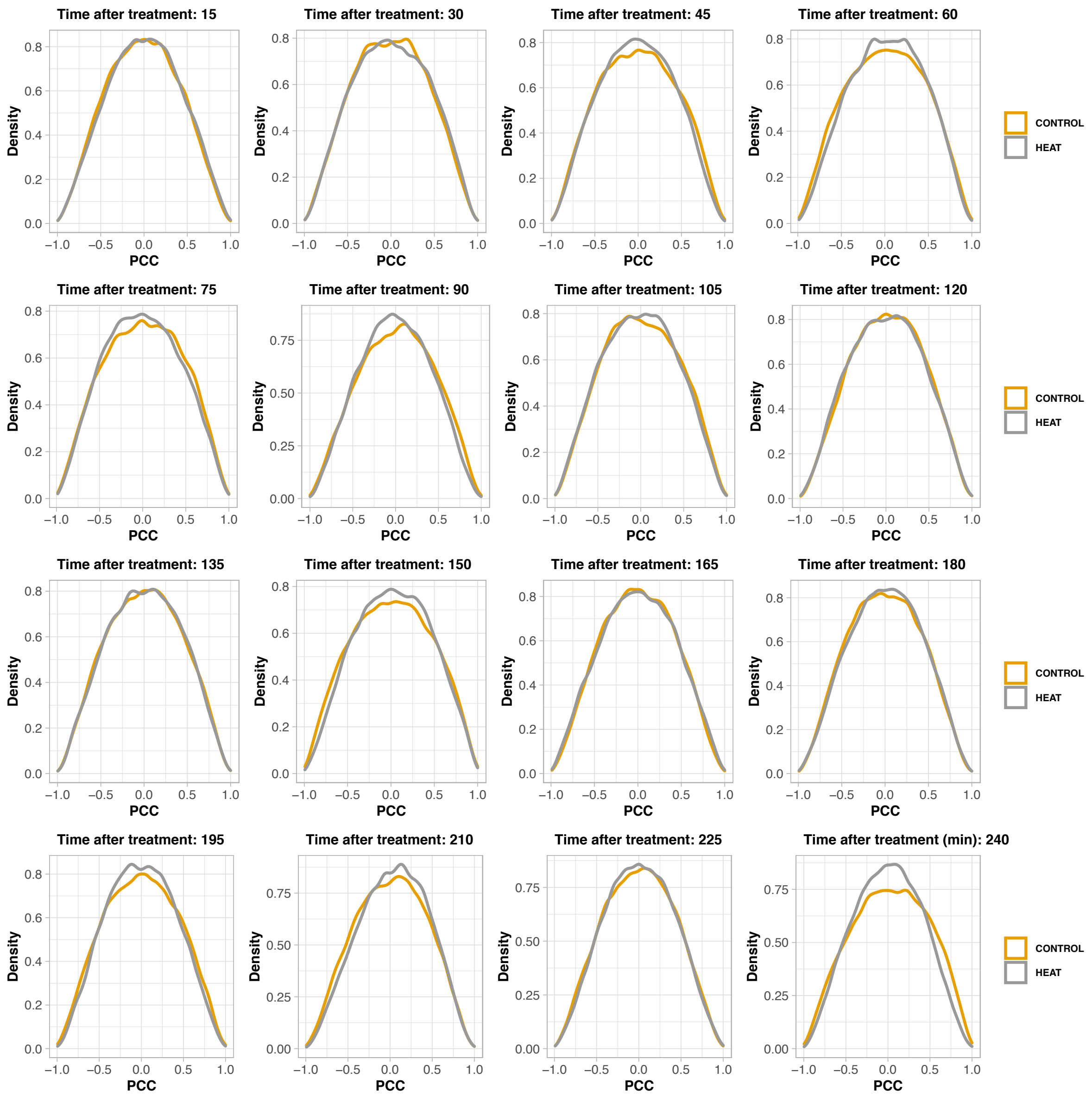


Figure S5: **Cross sectional population level correlation over the time course of heat stress treatment.** Each plot shows the distribution of Pearson Correlation Coefficient (PCC) at a given condition and time point (min) after the heat treatment.

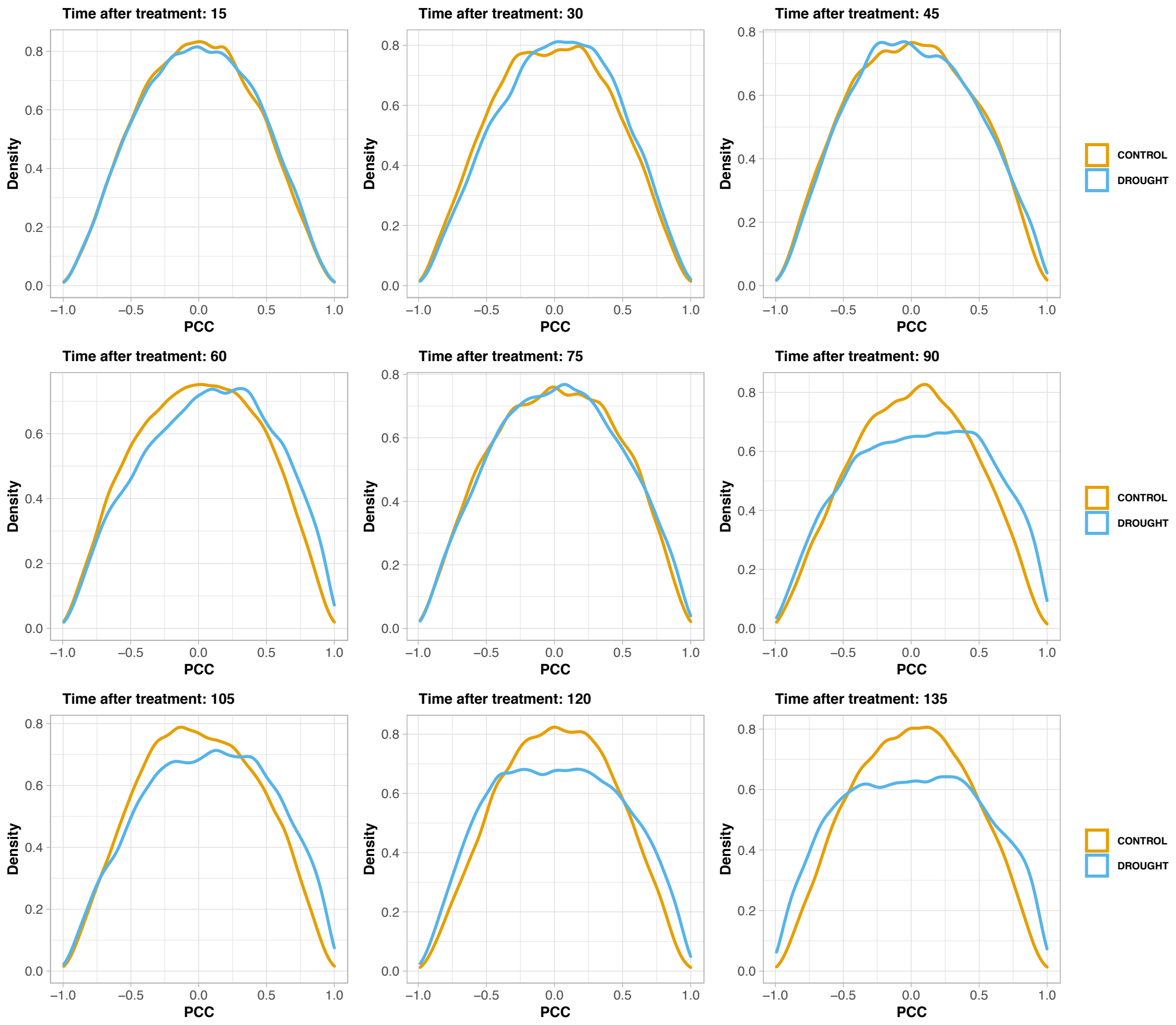


Figure S6: **Cross sectional population level correlation over the time course of drought stress treatment.** Each plot shows the distribution of Pearson Correlation Coefficients (PCC) at a given condition and time point (min) after the drought treatment.

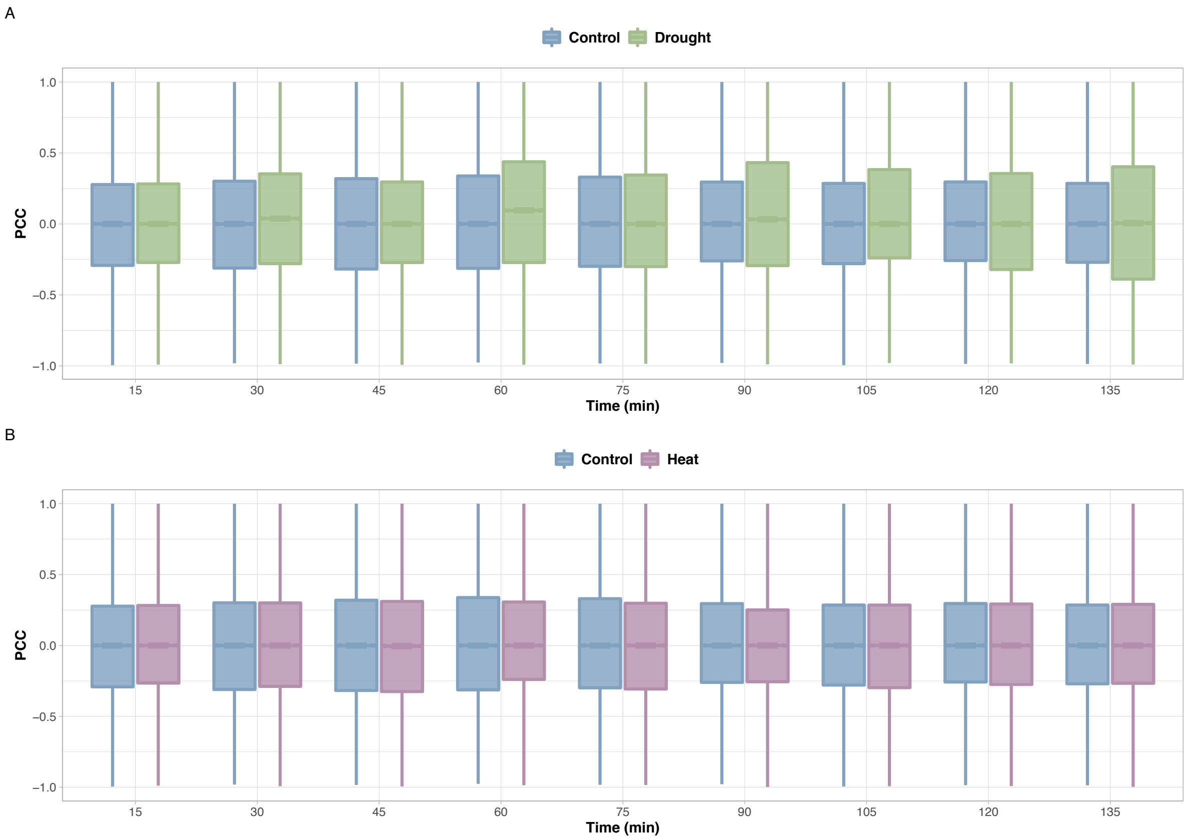


Figure S7: **Cross sectional population level Pearson Correlation Coefficient (PCC) show weak evidence of regulatory coherence.** Each boxplot shows the distribution of population correlation under a certain condition (control, drought, or heat) at a given time point.

#### C.3 Supplement figures for main Figure 4

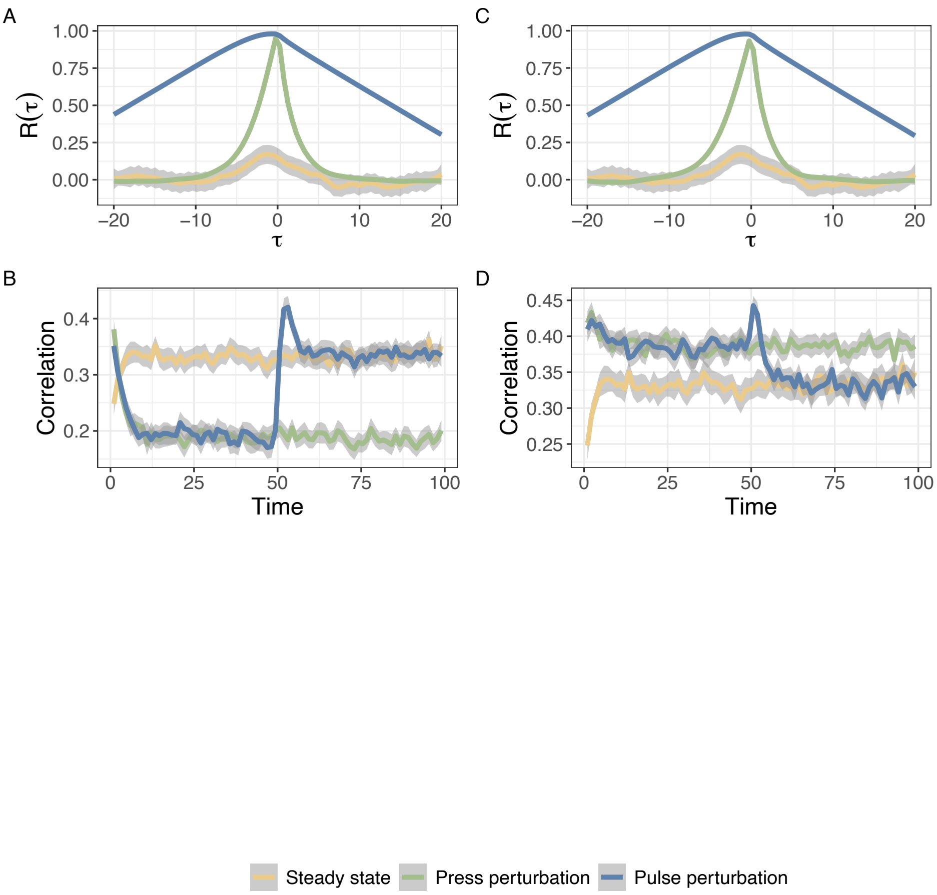


Figure S8: **Illustrated examples from stochastic simulation indicate the robustness of using temporal correlation to characterize the environmentally induced links.** Same parameter set of simulation with Fig. 2 but with 1000 population size (1000 cells). The cross correlation function and population-level correlation between activator $X$ and target $Y$ under a saturated regime **A and B**. The cross correlation function and population-level correlation between activator $X$ and target $Y$ under a saturated regime **C and D**. Colors represent three different types of external environmental conditions which lead to internal signaling (Steady state, press perturbation and pulse perturbation). $R_(\tau)$ is the cross correlation function with $\tau$ indicating the time delay. Note that the perturbation is imposed at $t = 0$ in **C** and **D**.

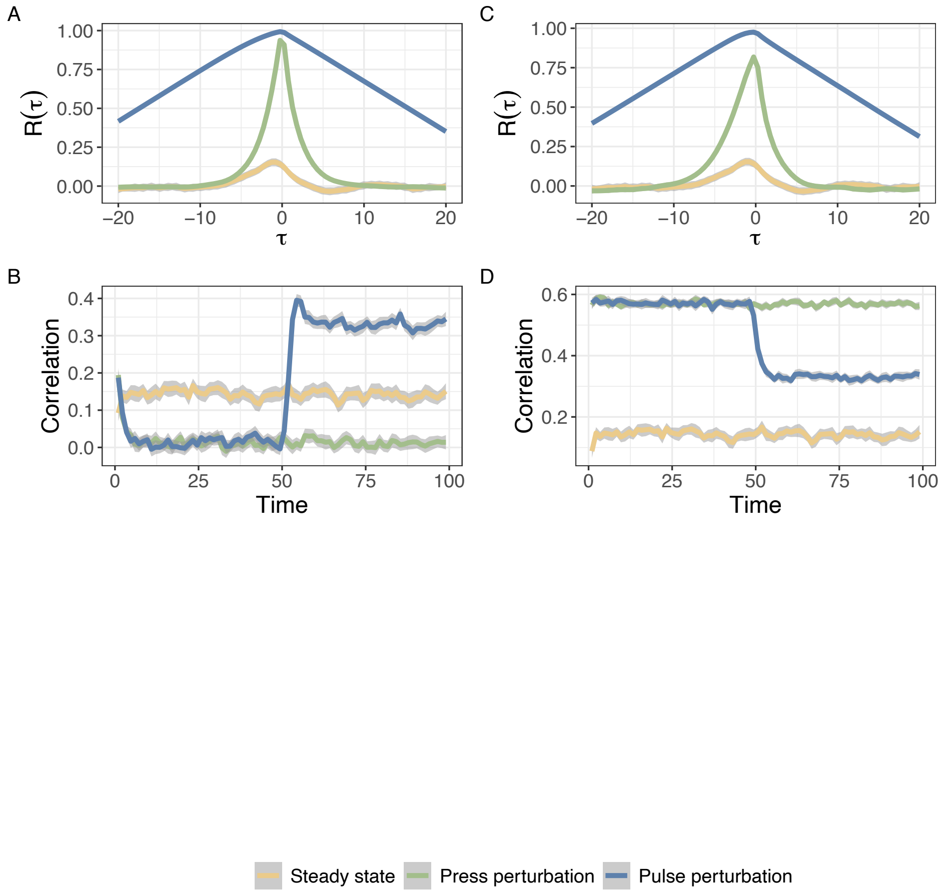


Figure S9: **Illustrated examples from stochastic simulation indicate the robustness of using temporal correlation to characterize the environmentally induced links.** The same plot with Fig. 2, but the perturbation is modulated through both binding affinity (K) and external signal. The cross correlation function and population-level correlation between activator $X$ and target $Y$ under a saturated regime **A and B**. The cross correlation function and population-level correlation between activator $X$ and target $Y$ under a saturated regime**C and D**. Colors represent three different types of external environmental conditions which lead to internal signaling (Steady state, press perturbation and pulse perturbation). $R_(\tau)$ is the cross correlation function with $\tau$ indicating the time delay. Note that the perturbation is imposed at $t = 0$ in **C** and **D**.

#### C.4 Supplement figures for main Figure 5

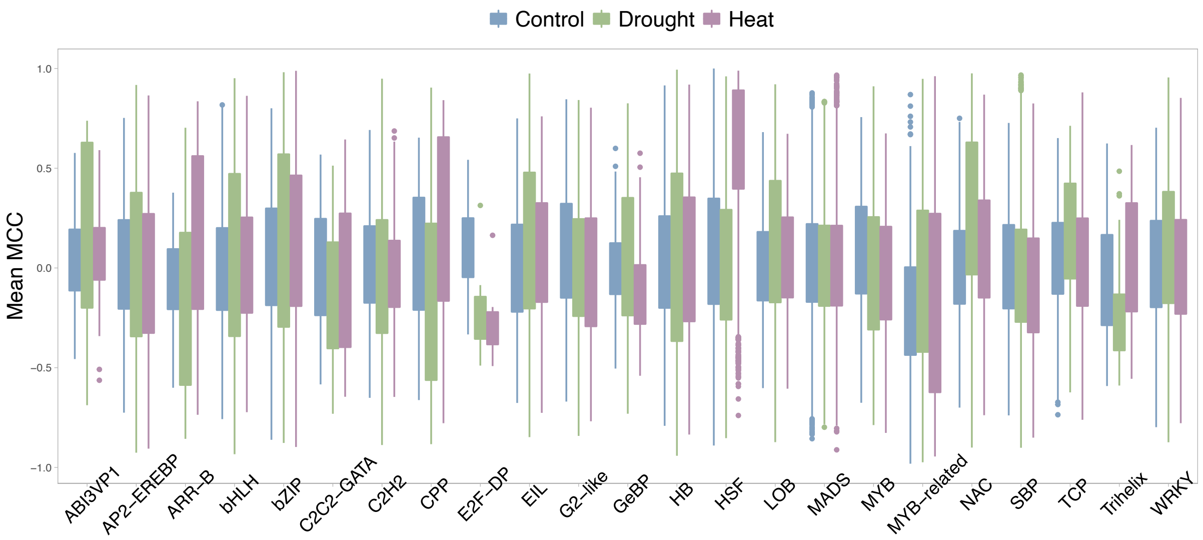


Figure S10: **Max Cross Correlation (MCC) for each regulator-target pair categorized by the regulator's family** The Max Cross Correlation (MCC) for each regulator-target pair in the network prior under multiple conditions, grouped by TF family.

#### C.5 Supplement figures for main Figure 6

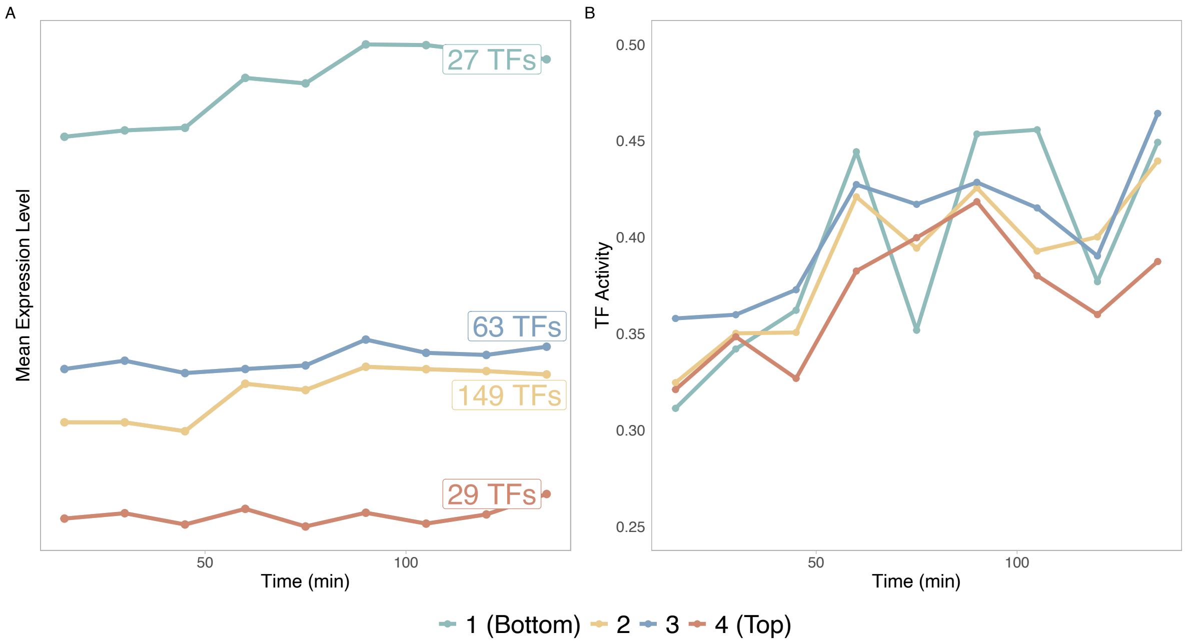


Figure S11: **Mean expression level and mean activity over time for TFs at each level in the hierarchy without removing all non-responsive TFs A.** Comparison of mean expression value of responsive TFs in the network hierarchy. The label of each line shows the number of TFs in that level. **B.** Dynamic TF activities calculated by average PCC of a TF with all of its target genes.

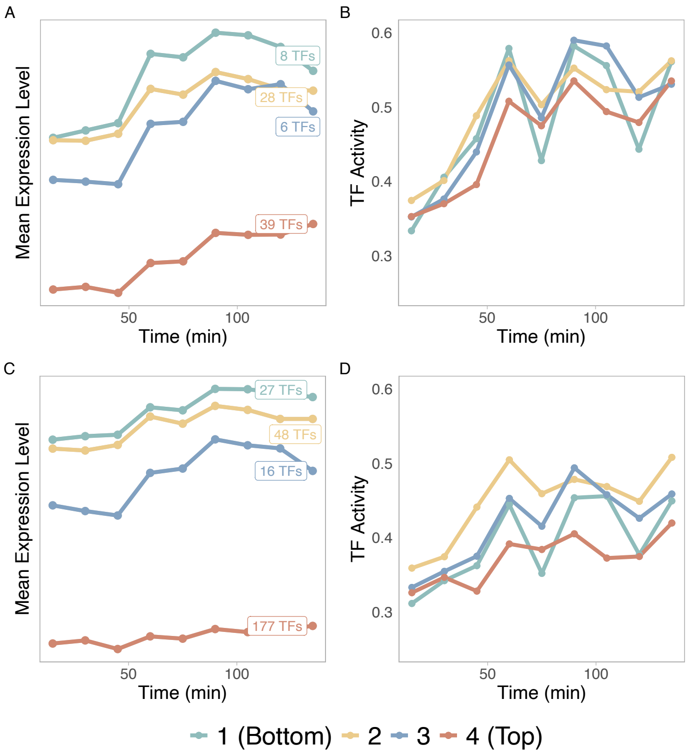


Figure S12: **Mean expression level and mean activity over time for TFs in the height-based hierarchy A.** Comparison of mean expression values of all responsive TFs in the network hierarchy with colors indicate the level of TFs. The label of each line shows the number of TFs within each level. **B.** Dynamic TF activities calculated by average PCC of a TF with all of its target genes. **C.** Comparison of mean expression value of all TFs (including all non-responsive TF) in the network hierarchy with colors indicate the level of TFs. The label of each line shows the number of TFs within each level. **D.** Dynamic TF activities calculated by average PCC of TFs at each level (including all non-responsive TFs).
